# Influence of pump laser fluence on ultrafast structural changes in myoglobin

**DOI:** 10.1101/2022.11.22.517513

**Authors:** Thomas R.M. Barends, Swarnendu Bhattacharyya, Alexander Gorel, Giorgio Schiro, Camila Bacellar, Claudio Cirelli, Jacques-Philippe Colletier, Lutz Foucar, Marie Luise Grünbein, Elisabeth Hartmann, Mario Hilpert, Philip J.M. Johnson, Marco Kloos, Gregor Knopp, Bogdan Marekha, Karol Nass, Gabriela Nass Kovacs, Dmitry Ozerov, Miriam Stricker, Martin Weik, R. Bruce Doak, Robert L. Shoeman, Christopher J. Milne, Miquel Huix-Rotllant, Marco Cammarata, Ilme Schlichting

**Affiliations:** Max Planck Institute for Medical Research, Heidelberg, Germany; Institut de Chimie Radicalaire, CNRS, Aix Marseille Univ, Marseille, France; Institut de Biologie Structurale, Grenoble, France; Paul Scherrer Institute, Villigen, Switzerland; European XFEL GmbH, Schenefeld, Germany; Laboratoire de Chimie, ENS de Lyon, Lyon, France; Department of Statistics, University of Oxford, Oxford, UK; ESRF, Grenoble, France

## Abstract

High-intensity femtosecond pulses from an X-ray free-electron laser enable pump probe experiments for investigating electronic and nuclear changes during light-induced reactions. On time scales ranging from femtoseconds to milliseconds and for a variety of biological systems, time-resolved serial femtosecond crystallography (TR-SFX) has provided detailed structural data for light-induced isomerization, breakage or formation of chemical bonds and electron transfer^1^. However, all ultra-fast TR-SFX studies to date have employed such high pump laser energies that several photons were nominally absorbed per chromophore^2-14^. As multiphoton absorption may force the protein response into nonphysiological pathways, it is of great concern^15^ whether this experimental approach^16^ allows valid inferences to be drawn vis-à-vis biologically relevant single-photon-induced reactions^17^. Here we describe ultrafast pump-probe SFX experiments on photodissociation of carboxymyoglobin, showing that different pump laser fluences yield markedly different results. In particular, the dynamics of structural changes and observed indicators of the mechanistically important coherent oscillations of the Fe-CO bond distance (predicted by recent quantum wavepacket dynamics^15^) are seen to depend strongly on pump laser energy. Our results confirm both the feasibility and necessity of performing TR-SFX pump probe experiments in the linear photoexcitation regime. We consider this to be a starting point for reassessing design and interpretation of ultrafast TR-SFX pump probe experiments^16^ such that biologically relevant insight emerges.

## MAIN

Light is an important environmental variable and organisms have evolved a variety of systems to sense it, exploit it, avoid it and deal with its damaging effects for example on DNA. Photosensory proteins contain a variety of light-absorbing chromophores with conjugated double bonds. Critical steps upon photon absorption include formation of a photoexcited chromophore, coupled electronically and vibrationally to the protein matrix, followed by transitions through a series of reaction intermediates. Elucidation of these events is not only of interest from a basic scientific point of view, but also of practical significance. Many photosensory proteins are either medically relevant (visual rhodopsins, melanopsins and cryptochromes), or useful tools for cell biology (imaging via fluorescent proteins, functional manipulations in optogenetics), or important for agriculture (photosystems and phytochromes). Of great interest to a very broad and large community is understanding of the relevant chemical mechanisms (including molecular determinants of quantum yields), the different photophysical and photochemical pathways and the origin of structural changes that accompany and effect biological function.

Until recently, experimental investigations of ultrafast events following photoexcitation were limited to various optical spectroscopies. Such studies provide deep insight into electronic and vibrational changes during the reaction but only restricted structural information, thereby limiting mechanistic insight. This shortcoming has been alleviated with the advent of X-ray free-electron lasers (XFELs), which provide highly intense short X-ray pulses that enable ultrafast time-resolved serial femtosecond crystallography (TR-SFX)^1^. Importantly, SFX allows the use of microcrystals. The high chromophore concentration in crystals results in high optical densities, which can be countered experimentally only by reducing crystal size. This is obligatory for efficient and well-defined initiation of photoexcitation reactions.

In time-resolved pump-probe SFX experiments, microcrystals are delivered into the XFEL beam using mostly liquid jets and diffraction data are collected at distinct time-delays following a photo-exciting pump laser flash. On the sub-ps to ns timescale, this approach has been used to study isomerization reactions in photoactive yellow protein (PYP)^3^, fluorescent protein^4^, various rhodopsins^5,6,8,9,11,13^ and phytochrome^7^; electron transfer reactions in a photosynthetic reaction center^10^ and photolyase^12^; photocarboxylation^17^ and photodissociation^2^. In all cases, a very high pump laser fluence was used to maximize the light-induced difference electron density signal, ^16^. As a result -when using the same cross sections for ground state and excited state absorption -significantly more than one photon is nominally absorbed per chromophore. Such excitation conditions differ markedly from those used in spectroscopic investigations, which are performed in the linear photoexcitation regime with generally much less than 0.5 photon/chromophore. Multiphoton artefacts are then avoided and only the biologically relevant single-photon reaction is probed. Consequently there can be considerable doubt as to whether SFX and spectroscopic measurements actually probe the same reaction, thus questioning the mechanistic relevance of the SFX results^18^. Nevertheless, the SFX community has failed so far to reach consensus on appropriate photoexcitation conditions for time-resolved pump probe experiments^16,19^.

Photodissociation of carboxymyoglobin (MbCO) is a well-characterized model reaction that has implications in a wide range of fields, ranging from organometallic chemistry to protein dynamics. The reaction has been studied by numerous computational and experimental approaches including TR-SFX^2^, with issues of high photoexcitation power density having been pointed out early on^20,21^. Here we examine the influence of the laser fluence on structural features of photoexcited MbCO derived from TR-SFX experiments. We show that the dynamics of structural changes differ and that indications for coherent oscillations of the Fe-CO bond distance predicted by recent quantum wavepacket dynamics^15^ are absent when using high photoexcitation power, which can be explained by the sequential absorption of two photons as inferred from quantum chemistry.

### Pump laser power titration

Power titration is a useful tool to establish the linear photoexcitation regime, namely that regime in which the magnitude of the response signal – or, in case of crystallographic investigations, the occupancy – increases linearly as a function of the incident laser energy density. Our first power titrations employed optical spectroscopy of the MbCO photodissociation reaction as a function of the power density of the pump laser. To this end, we determined the laser on -laser off difference absorption spectra (range 550-770 nm) 10 ps after photoexcitation by a 532 nm laser pulse. We explored different energy densities and pulse durations, specifically, three pulse durations of 80 fs, 230 fs and 430 fs at energy densities ranging from ∼1 to 90 mJ/cm² in the center of the Gaussian beam. The results are shown in Extended Data Fig. 1. The photolysis yield shows a clear dependence on the energy and duration of the pump pulse, with longer pulses being more efficient (up to ∼ 60% for the 430 fs pulse (Extended Data Fig. 1d)). At fluences above ∼20 mJ/cm², the shape of the transient difference spectra deviated from that of the static deoxyMb -MbCO difference spectrum, with a peak growing at ∼650 nm (Extended Data Fig. 1a-c1). Although this peak complicates estimation of the photolysis yield within the high energy density regime, it is clear that the linear photoexcitation regime lies below 10 mJ/cm^2^ (Extended Data Fig. 1d); this value might differ somewhat when photoexciting a microcrystal suspension.

To follow the CO photodissociation process at high temporal and spatial resolution, we performed a pump-probe TR-SFX experiment on MbCO at SwissFEL, yielding structures to 1.6 Å resolution (see Methods, Extended Data Table 1). The photolysis yield of Mb.CO microcrystals was determined using a laser power titration (laser fluence 6-101 mJ/cm^2^, see Extended Data Table 2 for excitation parameters) and performing TR-SFX at a 10 ps time delay. Inspection of 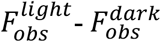 difference electron density maps shows a clear change of the magnitude of the peaks associated with bound and photolyzed CO, respectively, and the iron. At higher laser fluence, changes are also apparent in the protein and the porphyrin ring (Fig. 1a). Considering only the difference density as in previous TR-SFX studies^7,9,16,22,23^, a laser fluence of 101 mJ/cm^2^ appears preferable. However, further analysis of the data reveals that, for example, the fraction of photolyzed CO (denoted CO* henceforth) does not increase linearly at higher fluence, but instead levels off at ∼ 40 % (Fig. 1b). The underlying reason for the 40 % photolysis, despite very high laser fluence, is that a fraction of the thin plate-shaped MbCO crystals has at least one dimension that exceeds the 1/e laser penetration depth (∼ 7 µm), see Extended Data Fig. 2, Extended Data Table 2, Supplementary Note 1, Supplementary Fig. 1,2. Our previous investigation, using smaller crystals, showed 100 % photolysis^2^. Importantly, both observations demonstrate non-linear effects.

**Figure 1.**
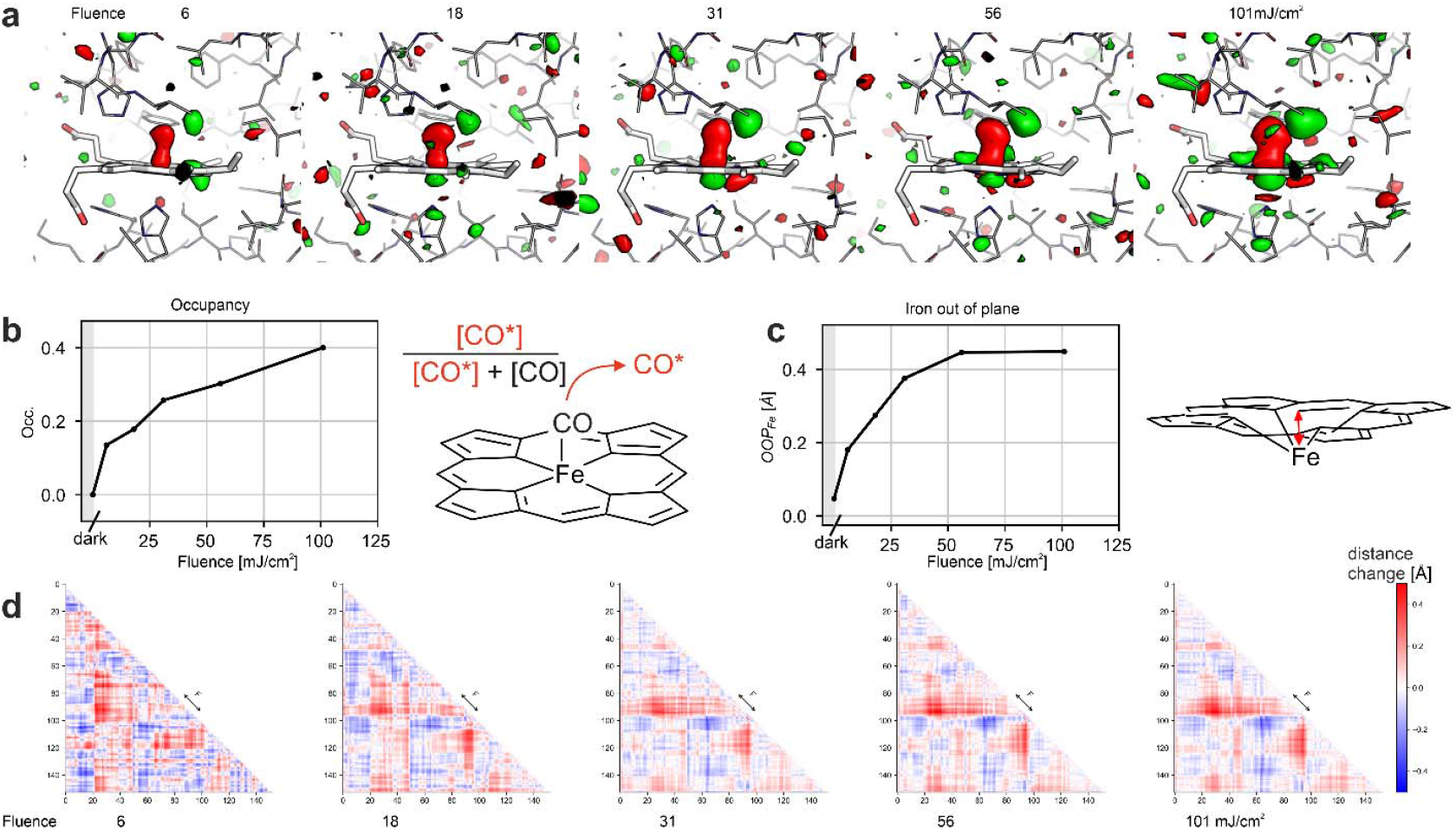
Crystallographic power titration at 10 ps time delay. a) Difference electron density maps, contoured at +3.0 (green) and -3.0 (red) sigma, overlaid on the dark-state structure of myoglobin for the various pump energies. b) Apparent occupancy of the CO* state as a function of pump laser fluence (mJ/cm^2^). The occupancy was determined by dividing the CO* peak height by the sum of the CO* and dark-state CO peak heights in mFo-DFc omit maps. c) Iron-out-of-plane distance as a function of pump energy. d) Cα-Cα-distance change matrices (“Go-plots”^67^) for the various pump laser fluences. Red indicates an increase, blue a decrease in distance. The F-helix (indicated) containing the heme-coordinating His93 moves away from several other elements (B, C, D, E, and G helices) and the E helix moves toward the FG corner and the H helix. Difference matrix plots between different pump laser fluences are shown in Extended Data Fig. 3.

**Figure 2.**
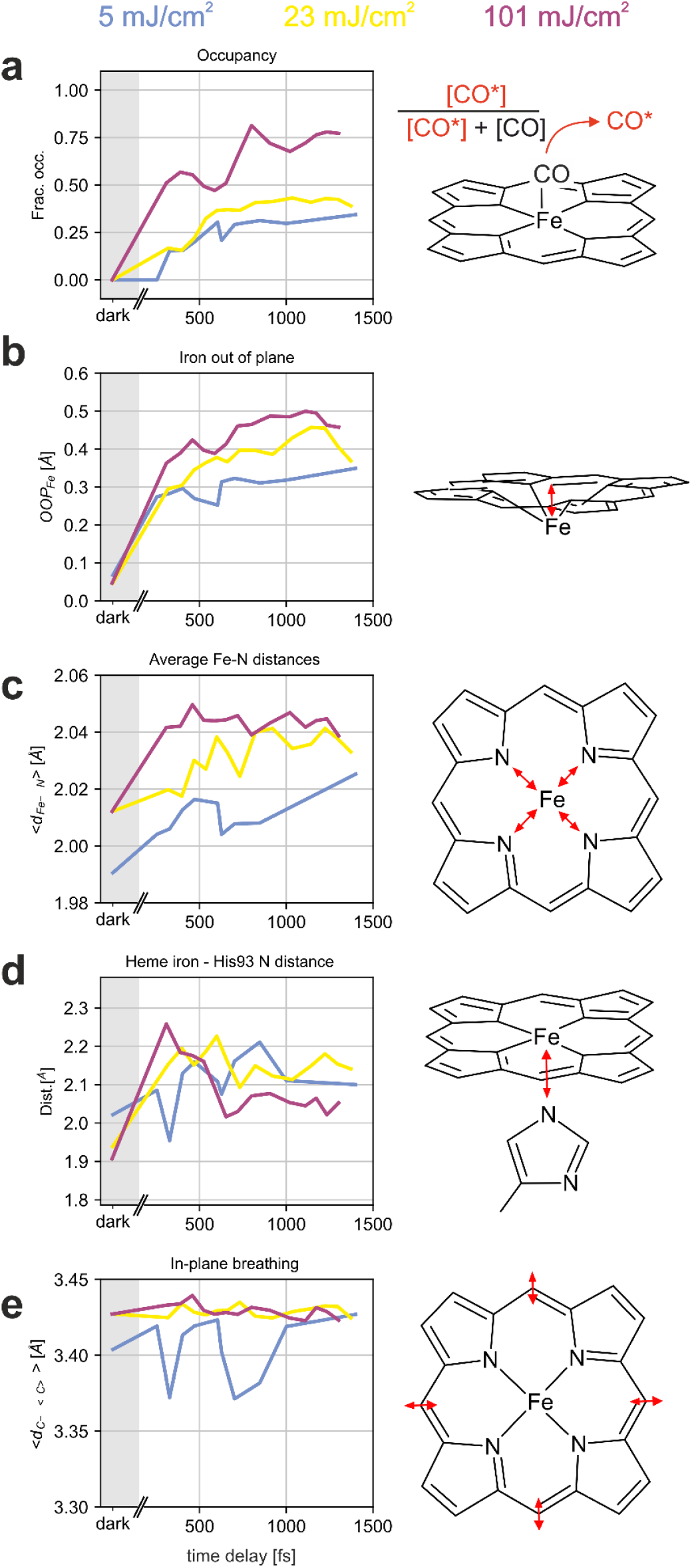
Heme structuraldynamics. a) Apparent CO*occupancy. Whereas at 5 and 23mJ/cm^2^ there is a smooth, slow increase, at 101 mJ/cm^2^ there is a rapid initial rise, followed by an equally slow increase to the final amplitude. The 101 mJ/cm^2^ curve can be understood as a super-position of contributions from the multiphoton-excited “front end” of the crystals with the few-photon excited “rear end” of the crystals(see Supplementary Note 1,Extended Data Fig. 2), resulting in almost instantaneous and apparently increasing occupancies of CO*, respectively. b) The iron-out-of-plane distance shows a larger amplitude with increasing fluence as does c) the average distance between the iron atom and the porphyrin N atoms. d) The distance between he me iron and proximalHis93 NE2 atom, too shows differences between the fluences used, with the lowest fluence showing an oscillation and the highest fluence first going up and then settling at a lower amplitude. e) The heme in-plane breathing (v7mode), determined as the average distance of the heme *meso* carbon atoms to the center of the heme, also varies with the fluence, with again the lowest fluence showing an oscillation and the higher fluences do not. Estimates for the oscillation periods are indicated by red dashed lines in Extended Data Fig. 5. That figure also shows co-ordinate uncertainties.

In the single photon excitation regime, increasing laser fluence raises the occupancy of the light-induced state, but does not affect the amplitude or nature of the structural or electronic changes. In addition to the nonlinear increase in CO* occupancy with laser fluence (Fig. 1b), the growing iron-out-of-plane distance (Fig. 1c) is a clear indication for nonlinear effects induced by multiphoton excitation. Although difference-distance matrix plots do not seem to show significant structural differences as a function laser fluence (Fig. 1d,Extended Data Fig. 3a), the analysis of the displacements of Cα atoms from the porphyrin nitrogen atoms indicates that such differences are indeed present (Extended Data Fig. 3b). Hence the influence of multiphoton excitation on structural changes it is not always immediately obvious and may demand very careful analysis.

### Structural changes at different fluences

To check whether the dynamics of the system are affected by the pump laser fluence, we performed TR-SFX at four different pump laser fluences (2.4, ∼5, 23, 101 mJ/cm^2^). These are within, higher but still within, outside, and far outside the linear excitation regime, respectively. To increase the relative yield of photoproduct at low laser fluence we used smaller crystals for the 2.4 and ∼5 mJ/cm^2^ data series (see Extended Data Table 2). The 2.4 mJ/cm^2^ data did not yield interpretable light-induced signal and will not be discussed further. The standard deviation of the time delays used in the SFX experiment is ∼100 fs for the 5, 23 and 101 mJ/cm^2^ data, taking into account timing jitter and the effects of data binning (see Methods section).

#### Dynamics of MbCO photolysis reaction

The hallmarks of MbCO photolysis are the observation of an unbound CO accompanied by changes in the iron’s spin states and position. Since CO photodissociates from the heme iron within 70 fs^24^, and in line with our previous TR-SFX experiment^2^ that showed full occupancy of CO* within the first time point, we anticipated no changes in CO* occupancy with time. Unexpectedly, however, our electron density maps show an apparent increase of the occupancy of CO* with time for the 5 and 23 mJ/cm^2^ data and, to a lesser extent of the 101 mJ/cm^2^ data (Fig. 2a, see below). Since the data series were collected during two beamtimes using different crystals, different dark state data and different laser settings (see Extended Data Table 2), it is very unlikely that this finding is a product of experimental errors. Instead, the ∼ 300 fs time constant of the apparent increase of CO* occupancy is reminiscent of the damping constant of a coherent nuclear oscillation of CO* that was predicted by recent computational wavepacket analysis^15^. Since the time resolution of our experiment does not allow the predicted 1 Å amplitude, ∼42 fs period oscillations to be resolved, they would manifest themselves simply as disorder due to distribution of the electron density over a large volume, resulting in an apparently low occupancy. As the oscillation damps, the CO* position “narrows” and its apparent occupancy converges to the value observed for the respective laser fluences at ∼10 ps (Fig. 1b). Importantly, the predicted CO* oscillation seems to be suppressed in the high photoexcitation regime; our previous high fluence study showed maximal CO* signal within the first time delay^2^. Similarly, at 101 mJ/cm^2^ - and in contrast to 5 and 23 mJ/cm^2^ data -we observe an initial rise to about 2/3 of the final value within the first time delay of our experiment, then the final 1/3 of the amplitude is reached with a speed comparable to what is observed at 5 and 23 mJ/cm^2^ (see Extended Data Fig. 2b).

We investigated the molecular basis for this observation by quantum chemical analysis. As described previously^15^, single photon absorption by MbCO results in wavepacket transfer from the ground state to the singlet Q state of porphyrin, followed by transfer to the singlet metal-to-ligand charge-transfer (MLCT) band. The wavepacket undergoes large-amplitude coherent oscillations in the Fe-CO coordinate on the singlet MLCT band. Importantly, strong Jahn-Teller distortions in the excited state affords an efficient energy transfer from the porphyrin plane (x,y-polarization) to the Fe-CO axis (z-polarization), activating dissociative stretching vibrations and thus CO dissociation^15^. To assess the quality of the quantum chemistry we computed the FeOOP distance using molecular dynamics in which a sudden dissociation of CO is imposed (see Supplementary Note 2). Since the results agree well with our SFX observations (Extended Data Fig. 4, Supplementary Note 2), we have high confidence in the accuracy of the computational approaches.

Our calculations show (see Supplementary Note 2) that in the high excitation regime the dissociation happens via a high-energy singlet state accessed by a sequential absorption of two photons. The first photon leads to the usual excited singlet Q state from which a second photon can be absorbed, as indicated by the absorption spectrum of the Q-excited heme-CO system (Extended Data Fig. 4). Analysis of the excitation character of this higher energy singlet state shows a mixed π→π* character of the heme and 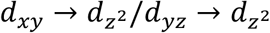 character with respect to the ground state, and therefore is dissociative for the Fe-CO bond (see Extended Data Fig. 4, Supplementary Note2). The potential energy surface (PES) of the singlet manifold along a relaxed scan coordinate at different fixed Fe-CO distances (see Methods, Computational Details section) clearly shows (Supplementary Fig. 3d) that, upon excitation to the dissociative singlet, after a second absorption from the Q-state, the excited wavepacket experiences a rapid decay towards Fe-C(O) dissociation. This dissociation is thus driven by the sudden change in electronic structure induced by photon absorption. Due to the (barrierless) repulsive nature of the potential, no coherent oscillations of the wavepacket are expected to be observed, in contrast to the single-photon regime, in which nuclear motions drive the electronic structural changes that lead to dissociation. This explains the quasi-instantaneous initial increase in apparent occupancy of CO* in our high fluence TR-SFX data. In conclusion, the photophysical mechanism of CO dissociation differs for single and two-photon absorption, respectively, resulting in different structural outcomes. This is in line with our experimental observations obtained under the respective photoexcitation conditions.

**Figure 3.**
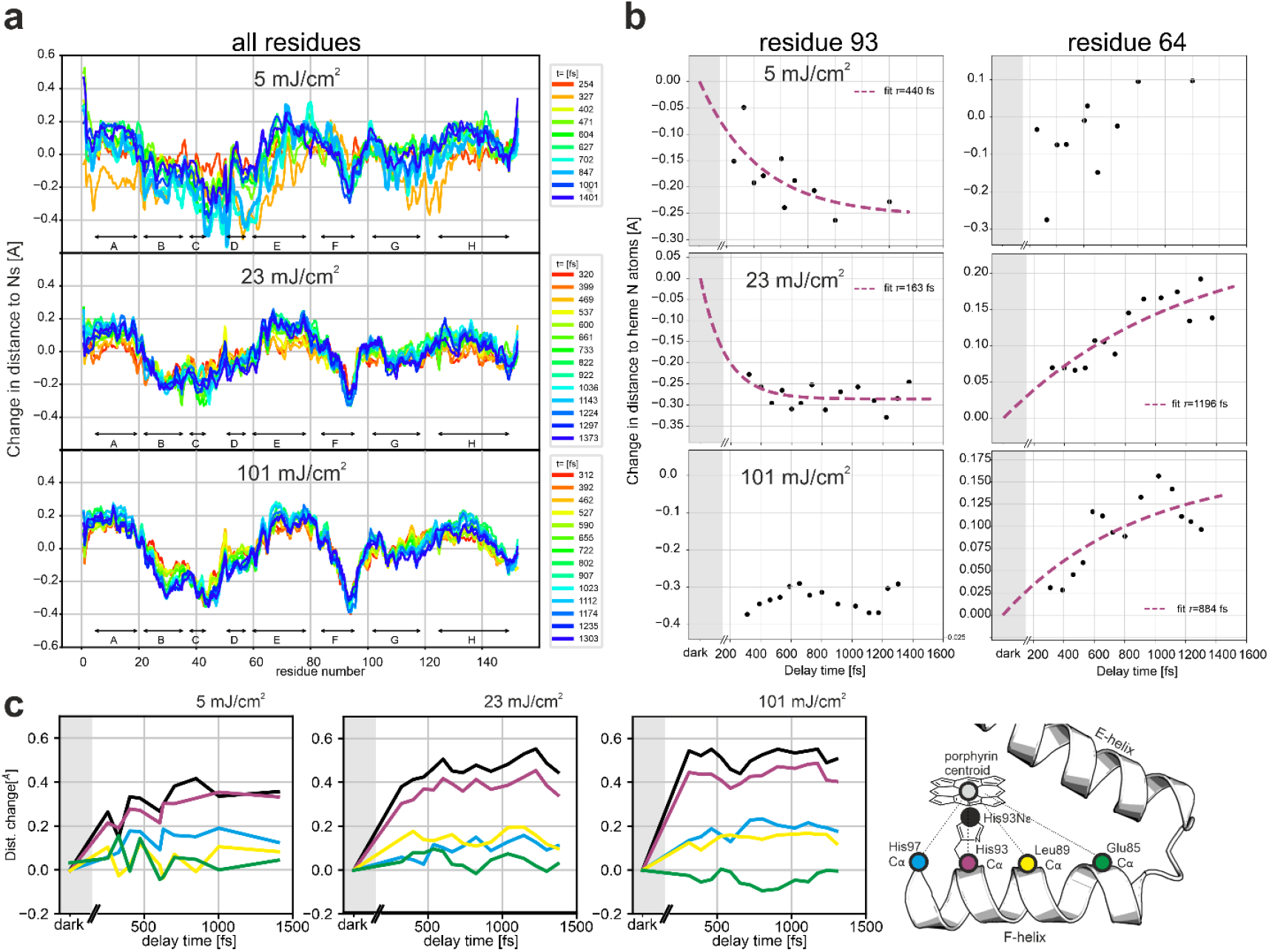
Dynamics of correlated structural dynamics upon MbCO photolysis depends on laser fluence. a) Guallar-type plots^29^, showing the change in distance of backbone N, Ca, and C atoms to the heme nitrogens for each time delay, for 5, 23, and 101 mJ/cm^2^ pump pulse energy. The speed of the changes is strongly fluence dependent. Correlated motions of helical elements show different temporal evolutions with time in particular for the 5 mJ/cm^2^ data, but move generally very fast in the 101 mJ/cm^2^ data, obscuring the sequence of events. For example, the displacement of the His93 main chain from the heme nitrogen atoms or heme centroid has a time constant of τ ∼ 320 fs and τ∼210 fs for the 5 and 23 mJ/cm^2^ data, respectively, but reaches its final value within the first time delay for the 101 mJ/cm^2^ data (b). In contrast, the movement of the distal His64 is hardly affected on the ultrafast time scale (b). The length of the correlated motion along the F-helix is clearly visible. Shown are displacements from the heme centroid of the His93 nitrogen (black) and Cα (red), the Cα atoms of His 97 (blue) and Glu85 (green) which are located at one helical turn upstream and downstream, respectively. Another turn further upstream the effect is strongly reduced. Importantly, a strong oscillatory modulation (period of ∼ 300 fs) is only visible for the 5 mJ/cm^2^ data.

#### Dynamics of the heme and coordinating His93

Upon CO photodissociation sequential changes of the Fe spin state occur, ultimately yielding the high spin (HS) state, and resulting in a movement of the iron out of the heme plane (FeOOP) as well as motions of surrounding protein moieties. Here, too, our observations show marked differences between the single- and multiphoton excitation regimes. The plot of the temporal evolution of the FeOOP distance shows a strong increase within the time-resolution of our experiment, resulting in ∼50% of the displacement, followed by a slower phase (τ ∼400 fs) as reported previously^2,15,25^, (Fig. 2b). In line with the observation at a 10 ps time delay, the FeOOP distance is largest for the 101 mJ/cm^2^ data. Upon Fe movement, the Fe distances to the nitrogen atoms of the pyrrole ring (Np) and of the proximal histidine (His93), respectively, increase (Fig. 2c,d). In the 101 mJ/cm^2^ data, the initially increasing Fe-His93 distance decreases again (Fig. 2d), in line with the larger FeOOP displacement (Fig. 2b) or due to increased vibrational energy redistribution^26^.

CO photodissociation also triggers heme breathing motions such as the ν7 in-plane vibration of the porphyrin ring^27^ which is predicted to have a distinct amplitude modulation with a period of ∼350 fs (see Fig. S12 in reference ^2^) due to the FeOOP movement. Although we cannot resolve the ν7 in-plane vibration itself (∼50 fs period), we do observe a ∼330 fs oscillation of the meso-carbon distances to the center of the heme for the 5 mJ/cm^2^ photoexcitation energy data (Fig. 2e); in contrast, the oscillation is hardly visible in the 23 and 101 mJ/cm^2^ data.

The heme dynamics have been studied by various spectroscopic methods, yielding time-constants of processes and proposals for the structural basis of the underlying molecular changes. Our structural data are in line with the interpretation of X-ray absorption spectroscopy data by Levantino et al^25^ proposing changes of the FeOOP distance, the Fe-Np and Fe-His bonds with a time constant of 70 fs, followed by a smaller change of the FeOOP distance with a time constant of 400 fs. The latter was suggested to be linked to a movement of the F-helix, which we, however, observe on a 200-300 fs time-scale depending on laser energy (see below). Our data do not agree with the structural interpretation by Shelby et al^28^, assigning a small FeOOP displacement to an 80 fs phase, followed by further FeOOP movement and elongation of Fe-Np bonds with a time constant of 890 fs.

#### Correlated protein structural changes

Oscillations of structural features (torsion angles, distances) of a light-sensitive cofactor and of near-by residues have been reported previously by TR-SFX^2,6^. These rapidly damped but coherent oscillations are a direct manifestation of the strong coupling of the chromophore and its environment. As in our previous study^2^, we observe oscillatory dynamics in the heme environment, reflecting coherent motions excited by photo-dissociation in the heme. In particular, the distal Hist93 χ2 rotation angle (Extended Data Fig. 6a) and the heme CMD to Lys42-O distance (Extended Data Fig. 6b) seem to mirror the modulation of the v7 oscillation (Fig. 3e) as does the Ser92-His93 hydrogen bond length (Extended Data Fig. 6c). The χ2 torsion angle of Phe43 (Extended Data Fig 6d) and the heme CHD-Ile99 CD1 distance (Extended Data Fig. 6e), on the other hand, appears to mirror the heme doming frequency, see also ref. ^2^. Importantly, the temporal development of these angles and distances show marked differences between the low- and high fluence regimes (Extended Data Fig. 6, 7).

Sequence displacement graphs^2,29^ —which illustrate the change in distance of the protein main-chain atoms to the center of the four porphyrin N atoms as a function of the time delay between the pump and probe pulses— show substantial main-chain changes within 1 ps throughout the whole protein for all pump laser fluences, but the dynamics differ dramatically (Fig. 3a,b). For many structural elements, the 5 and 23 mJ/cm^2^ data display a temporal evolution over the entire ultrafast time-series -whereas the 101 mJ/cm^2^ data show the essentially the entire displacement within the first time point, similar our previous observation (Suppl. Fig S5a)^2^. This is particularly noticeable for the displacement of the proximal His93 from the heme and the coupled motion of adjacent residues (Fig. 3). Moreover, a strong oscillatory modulation with a frequency of ∼ 300 fs of the His93 displacement and the neighboring residues (Fig. 3c) is clearly visible for the 5 mJ/cm^2^ data only. Thus, the multiphoton effects are not limited to the small-scale motions of a few atoms but also affect larger-scale correlated protein motions in the entire protein (Extended Data Fig. 8), including the radius of gyration Rg (Extended Data Fig. 8). As for other displacements, the oscillations are pronounced in single photon excitation data (5 mJ/cm^2^ fluence).

The striking change in dynamics of correlated motions (Fig. 3) with laser fluence is likely due to the excess energy deposited in the heme and Raman-active modes via multiphoton absorption, ultimately resulting in heating^21^. At higher temperature, the displacement of the atoms from their equilibrium position increases so that modes sample more of the anharmonic part of the potential energy surface. As the rate of energy transfer between modes depends on the nonlinear coupling between them^30^, they are then in effect more strongly coupled^21^, resulting in faster structural changes.

## Conclusions

The combination of spectroscopy, TR-SFX and quantum chemistry provides unprecedented insight into reaction mechanisms and protein dynamics, in particular when the initial ultrafast steps can be analyzed as fully as only light-triggered reactions allow. An implicit assumption in such studies is that all three approaches study the same reaction, namely one triggered by the absorption of a single photon. Hence, photoexcitation conditions matter. Recent quantum dynamics computations have linked the microscopic origins of ligand photolysis and spin crossover reactions in photoexcited MbCO to nuclear vibrations and predicted coherent oscillations of the Fe-CO bond distance^15^. This prediction is consistent with our TR-SRX data showing an apparent increase of the CO* occupancy within 0.5 ps after low fluence photoexitation of MbCO, which mirrors the damping of the oscillation. In addition to providing first experimental support of this computational prediction, our low fluence TR-SFX data also allow correlating of spectroscopically derived information^25,28^ with structural data, including the coupling of modes^31^. Although our time resolution does not allow observation of the predicted coupling of the heme doming mode and the 220 cm^-1^ (150 fs period) Fe-His mode^31^, we observe the coupling of the FeOOP mode and the in plane heme breathing mode.

High fluence excitation results in multiphoton absorption in MbCO. Our computations show that sequential two-photon excitation changes the photophysical mechanism by directly populating a dissociative state, bypassing the wavepacket oscillations, and thus explain the distinct TR-SFX results under high fluence photoexcitation conditions. Moreover, multiphoton excitation results in the deposition of excess energy into the system, which opens further relaxation pathways because the thermal decay channel is strongly coupled to collective modes of protein^21,32^. It was shown previously^21^ that under high excitation conditions MbCO displays power-dependent features with subpicosecond components attributed to increased anharmonic coupling between the collective modes of the protein and the increased spatial dispersion of the larger amount of excess energy. Indeed, we observe faster and larger structural changes when using high fluence photoexcitation outside the linear regime (Fig. 3a, c). The changes are not purely isotropic but correlate with the energy flow, for example, the F-helix -which is directly linked to the heme via the proximal His93 -is much more affected than the distal E-helix containing His64 (Fig. 3b). Moreover, the influence of the photoexcitation regime on oscillatory motions – which are much more pronounced in the low fluence data (5 mJ/cm^2^) -complicates identification of coherent oscillations that are involved in mode coupling and ultimately result in the biologically relevant structural changes.

Given the widespread and continuing^16^ use of overly high photoexcitation energies, it is highly likely that the light-induced structural changes described for other systems also involve multiphoton effects but that were presented and interpreted as mechanistically relevant. Likely symptoms include large structural changes on the ultrafast time-scale^7,10,13^, including those referred to as protein quakes^5,33^ and conformational transitions that are not in line with spectroscopic results^7,34^. Our results call into question recent statements promulgating the value of TR-SFX pump-probe experiments performed above single-photon excitation thresholds^16^.

## Methods

### Sample preparation

Horse heart myoglobin (hhMb) was purchased from Sigma Aldrich (M1882). After dissolving lyophilized hhMb powder (70 mg/ml) in 0.1 M Tris HCl pH 8.0, the solution was degassed and then treated with CO. Upon addition of sodium dithionite (12 mg/ml) while constantly bubbling with CO gas, the color of solution turned to raspberry red. Dithionite was removed by desalting the protein solution via a PD10 column equilibrated with CO saturated 0.1M Tris HCl pH 8.0. Subsequently, the MbCO solution was concentrated to ∼ 6 mM using centrifugal filters before freezing in liquid nitrogen for storage.

hhMb crystals were grown in seeded batch by adding solid ammonium sulfate to a solution of 60 mg/mM hhMB in 100 mM Tris HCL pH 8.0 until the protein started to precipitate (∼3.1 M NH_3_SO_4_). Seed stock solution was then added. Crystals appeared overnight and continued growing for about a week, yielding relatively large, often intergrown plate-shaped crystals^2^. Using an HPLC pump the crystalline slurry was fractured using tandem array stainless steel ¼inch diameter filters^35^. For beamtime 1 (March experiment) the first tandem array contained 100 and 40 µm filters followed by a second tandem array of 40, 20, 10 and 10 µm filters. For beamtime 2 (May experiment), the crystals were further fractured using a tandem array of 10, 5, 2 and 2 µm stainless steel ¼inch diameter filters. On average, the largest crystal dimensions of the crystallites were ∼15 µm (Supplementary Fig. 1a) and ∼9 µm (Supplementary Fig. 1b) for beamtimes 1 and 2, respectively.

### Laser power titration

Time-resolved spectroscopic data for estimating the extent of photolysis as a function of laser power density were obtained using a 6 mM hhMbCO solution. The sample was placed in a rectangular borosilicate glass tube sealed with wax to keep the solution CO saturated. The optical path length was 50 μm and the thickness of the glass tube was 1mm. The optical density at the pump laser wavelength (532 nm) was ∼0.5. An identical tube filled with the buffer solution (0.1 M Tris HCl pH 8.0) was used as a blank.

The fs laser pulses were generated by a Ti-sapphire amplifier (Legend, Coherent) seeded by a Mira fs oscillator. The laser output was divided into two branches: The vast majority was used as input of an optical parameteric amplifier (Topas, LightConversion) to generate the pump pulses at 532 nm, while the remaining fraction was sent onto a sapphire crystal to generate short white-light pulses. Correction for white light temporal chirp (of <2ps over the probed window) was not needed at the time delay of interest. Mechanical choppers were used to lower the original 1 kHz repetition rate of both pump and probe pulses to 1 Hz and 500 Hz respectively. Pump and probe beams were spatially and temporally overlapped at the sample position and the relative time delay was set using a delay line. Pump pulses were focused to a FWHM of about 0.1 mm, while the probing white-light FWHM beam size was about 0.02 mm diameter (FWHM). Each time-resolved spectrum was obtained by averaging 60 consecutive pump-probe events. A Berek compensator was used to change the pump light polarization from linear to circular. The 80 fs pump pulses were stretched to ∼230 fs and ∼430 fs by inserting 10 and 20 cm water columns, respectively, along the pump laser path^36^. The difference spectra shown in Extended Data Fig. 1 were obtained using linearly polarized pump light; analogous results were found using circularly polarized light (data not shown).

### Data collection at SwissFEL

The TR-SFX experiment was performed in March (beamtime 1)/May (beamtime 2) 2019 using the Alvra Prime instrument at SwissFEL^37^ (proposal #20181741). To follow the time-dependent light–induced dynamics, an optical pump, X-ray probe scheme was used. The repetition rate of the X-ray pulses was 50 Hz. Diffraction images were acquired at 50 Hz with a Jungfrau 16M detector operating in 4M mode. The outer panels were excluded to reduce the amount of data.

The X-ray pulses had a photon energy of 12 keV and a pulse energy of ∼500 μJ. The X-ray spot size, focused by Kirkpatrick–Baez mirrors, was 4.9 × 6.4 μm^2^ in March 2019 and 3.9 × 4.1 μm^2^ in May 2019 (horiz. × vert., FWHM). To reduce X-ray scattering, a beamstop was employed and the air in the sample chamber was pumped down to 100–200 mbar and substituted with helium. The protein crystals were introduced into the XFEL beam in a thin jet using a gas dynamic virtual (GDVN) nozzle injector^38^. The position of the sample jet was continuously adjusted to maximize the hit rate. In the interaction point, the XFEL beam intersected with a circularly polarized optical pump beam originating from an optical parametric amplifier producing laser pulses with 60 ± 5 fs duration (FWHM) and 530 ± 9 nm (FWHM) wavelength focal spots of 120 × 130 μm^2^ and 150 × 120 μm^2^ (horiz. × vert., FWHM), in March and May, respectively. The laser energy was 0.5 and 1 μJ in May and 1-18 μJ in March 2019, corresponding to laser fluences of ∼2.5 to ∼101 mJ/cm^2^ and laser power densities of ∼40 to 1700 GW/cm^2^ (see Extended Data Table 2). Using an absorption coefficient of 11,600 M^-1^cm^-1^ for horse heart carboxymyoglobin at 530 nm, this results in nominally ∼0.3 to 12 absorbed photons/heme at the front of a crystal facing the pump laser beam. Time-zero was determined in the pumped-down chamber at the same low-pressure helium atmosphere used for data collection. Information from a THz timing tool was used for determining the actual time delay. A power titration was performed at a 10 ps time delay (March 2019). Full time series were collected for pump laser fluences of 5 (May), 23 and 101 mJ/cm^2^ (March). For the 5 mJ/cm^2^ time series, the time delay could be set with sufficient reproducibility that each time point could be collected as a single data set, with nominal time delays of Δ*t* = 150, 225, 300, 375, 450, 525, 600, 750, 900, and 1300 fs. Using the timing tool available at the beam line, the actual time delays of these datasets could then be determined to be 254, 327, 402, 471, 627, 702, 847, 1001 and 1401 fs, with widths of ∼85 fs. The number of images in each data set ranged from ∼10,000 to >30,000, with >60,000 in the dark data set. At the time the 23 and 101 mJ/cm^2^ time series were collected, the available timing reproducibility was less, and data sets were collected at a series of preset nominal time delays ranging from 150 to 1300 fs that were then merged into large sets of ∼150,000 images for both fluences. These where then sorted according to the actual time delay of each image as determined by the timing tool of the beam line. Then, the data were split into smaller datasets by moving a window of 20,000 images over the data for each fluence in steps of 10,000 images. The timing distributions of these partial datasets have standard deviations of between 40 and 70 fs. In combination with the accuracy of the timing tool we estimate the true widths of these distributions to be ∼100 fs. It should be noted that the overlap of the time delay distributions caused by this “binning” of the 23 and 101 mJ/cm^2^ data will result in a “smearing out” of time-dependent effects. In each case, every 11^th^ pulse of the pump laser was blocked, so that a series of ten light activated and one dark diffraction pattern were collected in sequence. High-quality dark data sets were generated by merging all laser-off patterns as well as separately collected, dedicated laser-off runs.

The latter were also used to confirm that the interleaved dark data in the light runs were indeed dark and not illuminated accidentally.

### Diffraction data analysis

Diffraction data were processed using CrystFEL 0.8.0^39^; Bragg peaks were identified using the peakfinder8 algorithm and indexing was performed using XGANDALF^40^, DIRAX^41^, XDS^42^ and MOSFLM^43^. After Monte-Carlo integration^44,45^, occupancies of the photolyzed state were determined by calculating 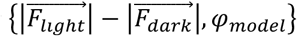 electron density maps using phases from a model without the CO ligand. The heights of the peaks for the CO in the ground (dark) and photolyzed CO* states were then used to calculate the occupancy *f* using:

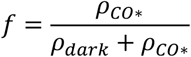

where *ρ*_*CO∗*_ and *ρ*_*dark*_ are the peak heights for the dark- and CO*-state CO peaks, respectively. These occupancies are shown in Figures 1 and 2.

To obtain refined structures of the photolyzed states, structure factors were extrapolated to full occupancy using the linear extrapolation approximation^46,47^. Briefly, the amplitudes

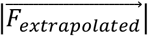 corresponding to 100% occupancy were calculated using the formula:

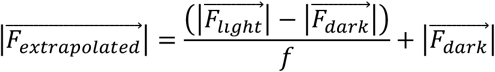

where 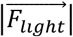 and 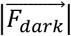 are the measured amplitudes of the light- and dark state structure factors, respectively, and *f* is the estimated occupancy of the photolyzed state.

Importantly, we found that the best apparent occupancy to be used for extrapolation (that is, the occupancy that results in maps that exclusively show the photolyzed state) differs from those found from difference electron density maps and is sensitive to resolution limits, weighting schemes etc. and must be determined a new for each case. This was done by increasing the assumed occupancy until dark state features became apparent in the extrapolated electron density maps^46-48^, i.e. where density for unphotolyzed (i.e. heme-bound) CO became visible. To this end, light data were scaled to the dark data using SCALEIT^49^ from the CCP4 suite^50^ using Wilson scaling. After scaling, light-dark differences were calculated and used for the calculation of extrapolated structure factors, using assumed occupancies ranging from 0.05 to 0.7 in steps of 0.05. Each set of extrapolated amplitudes was combined with dark state phases and an 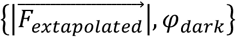 map was calculated using PHENIX^51^. The electron density in this map at the position of the dark-state CO oxygen atom was determined and plotted against the assumed occupancy. The resulting data points were fitted with a simple asymptotic function, and the occupancy at which this function crossed 1.0 σ (i.e. where an unphotolyzed CO would have become visible) should then yield the occupancy for a particular time delay.

These apparent occupancies, too, showed an asymptotic increase with delay time as did those determined from difference map peaks (see above). As we interpret this to be an artefact caused by disorder of the CO at early time delays, the final occupancies used to calculate extrapolated structure factors were set to the plateau value estimated for the respective pump pulse fluences (0.2, 0.3 and 0.42, for 5, 23 and 101 mJ/cm^2^ fluence, respectively). A model of photolyzed CO myoglobin was then refined against each of the resulting extrapolated data sets using phenix.refine build 1.19.2_4158^45^, using a heme geometry in which the planarity restraints were relaxed to allow the heme to respond to photolysis. We investigated the use of different low- and high-resolution limits. Using a low-resolution limit of 30 Å worked for some data sets, but for others resulted in problems during light-dark scaling, likely due to differences in beam stop placement. However, we found that a low-resolution limit of 10.0 Å could be used for all data sets and therefore imposed this for all structure factor extrapolations. Moreover, while the individual dark- and light datasets extend to 1.4-1.3 Å resolution, paired refinement^52^ suggested that the extrapolated data were useful in refinement to a resolution of 1.6 Å (not shown). This is due to the errors introduced by the extrapolation process^46-48^. Indeed, extrapolation may even introduce negative structure factor amplitudes into the data set. However, at 1.6 Å resolution less than ten percent of our extrapolated structure factors are affected by this problem and “rescuing” them using the methods recently evaluated by de Zitter and coworkers^53^ did not result in appreciable improvements. We therefore did not apply these methods. The dark data were used to their respective resolution limits. We also evaluated the usefulness of Q-weighting^54^ in the calculation of extrapolated structure factors, but found no apparent improvement in data quality and therefore did not use it for the results presented here.

Structures were analyzed using COOT^55,56^, PYMOL^57^ and custom-written python scripts using NumPy^58^ and SciPy^59^. To obtain error estimates for structural parameters such as bond lengths and torsion angles, bootstrap resampling was performed as follows: of each dataset, ∼100 resampled versions were created using a sample-and-replace algorithm. These were used to refine ∼100 versions of each structure, which were used to determine standard deviations. The number of 100 resampled versions was chosen as this has been shown to result in sufficient sampling^48,60^ while still being computationally tractable.

### Quantum Chemistry

For the calculation of the absorption spectra and attachment-detachment density analysis, a reduced model in gas phase was constructed that includes the Fe-porphyrin along with CO on one side of the porphyrin plane and an imidazole (part of the proximal histidine) on the other side. The geometry was optimized at the DFT/B3LYP/LANL2DZ level. The absorption spectra were computed at the optimized singlet ground state geometry at XMS-CASPT2/CASSCF/ANO-RCC-VDZP level using OpenMolcas^61,62^. An active space of 10 electrons in 9 orbitals was used (5d orbitals of iron and 4 π orbitals). The stick spectra were convoluted with Gaussians of 0.1 eV full width at half maximum to obtain the spectral envelope.

For the relaxed scan along the Fe-C(O) dissociation coordinate, the geometries of the model system were optimized at fixed Fe-C(O) bond lengths on the lowest quintet ground state at the DFT level. XMS-CASPT2 calculations were performed at these geometries to obtain the PES cut, to extract 60 singlets included in the state-averaging to account for the dissociative state corresponding to the sequential two-photon absorption model.

The QM/MM model was constructed on the basis of the crystal structure of the horse heart myoglobin (PDB code 1DWR)^63^. The protein was solvated in a box of 11684 water molecules. First, a minimization of the whole system was performed, followed by an NVT dynamics of 125 ps and a production run of 10 ns using Tinker 8.2.1^64^. From the MD, we extracted several snapshots to perform QM/MM MD, using a development version of GAMESS-US/Tinker^65^ The QM region includes the heme, CO and parts of the proximal and the distal histidines and was described at the DFT level. The rest of the system is described at the MM level with the CHARMM36m^66^ force field. A time step of 1 fs was used for the QM/MM molecular dynamics simulations.

## Acknowledgements

We are grateful to Matteo Levantino for providing UV-vis reference spectra of deoxy myoglobin and Mb.CO. SB, MHR and MC acknowledge the support by the French National Research Agency via the grant ANR-19-CE29-0018 (MULTICROSS). We acknowledge support by the Max Planck Society.

## Data Availability and Code Availability

Structures have been deposited with the PDB (accession codes 8BKI, 8BKJ, 8BKK, 8BKL, 8BKM, 8BKN, 8BKH, 8BKO, 8BKP, 8BKQ, 8BKR, 8BKS, 8BKT, 8BKU, 8BKV, 8BKW, 8BKX, 8BM8, 8BMA, 8BMB, 8BMC, 8BME, 8BMF, 8BMG, 8BMH, 8BMI, 8BMJ, 8BMK, 8BML, 8BMM, 8BMN, 8BNC, 8BND, 8BNE, 8BNF, 8BNG, 8BNH, 8BNI, 8BNJ, 8BNK, 8BNL, 8BNM, 8BNN, 8BNO, and 8BNP), stream files, extrapolated structure factor amplitudes, analysis scripts and relaxed heme geometry description with zenodo.com under doi 10.5281/zenodo.7341458.

Analysis scripts can be retrieved from https://github.com/tbarends/

## Supplementary Information is available for this paper

Correspondence and requests for materials should be addressed to TRMB, MHR, IS.

## Author Contributions

E.H., R.L.S., I.S. prepared sample, G.S, M.C. performed optical power titration, P.J.M.J., G.S, M.C., did laser work at SwissFEL,

G.S., C.C., P.J.M. J., G.K., C.J.M., M.C. operated the Alvra instrument at SwissFEL; T.R.M.B, A.G., G.S., C.C., J-P.C, L.F.,M.L.G.,M.H., P.J.M.J., M.K. G.K. K.N. G.N.K., D.O.M.S. M.W., R.B.D. R.L.S, C.J.M. I.S. performed data collection at SwissFEL;

M.K., M.L.G, M.S., G.N.K., R.L.S., R.B.D. injected the crystals at SwissFEL;

T.R.M.B, A.G., J-P.C, L.F., M.H., K.N., D.O. performed data analysis at SwissFEL;

S.B. and M.H-R. performed quantum chemistry calculations;

T.R.M.B., A.G. and M.H. performed off-line data analysis; data were analyzed by T.R.M.B. and I.S.; C.B. and B.M. provided spectroscopic input; T.R.M.B and I.S. wrote the manuscript with input from all authors.

## Extended Data

**Extended Data Table 1:**
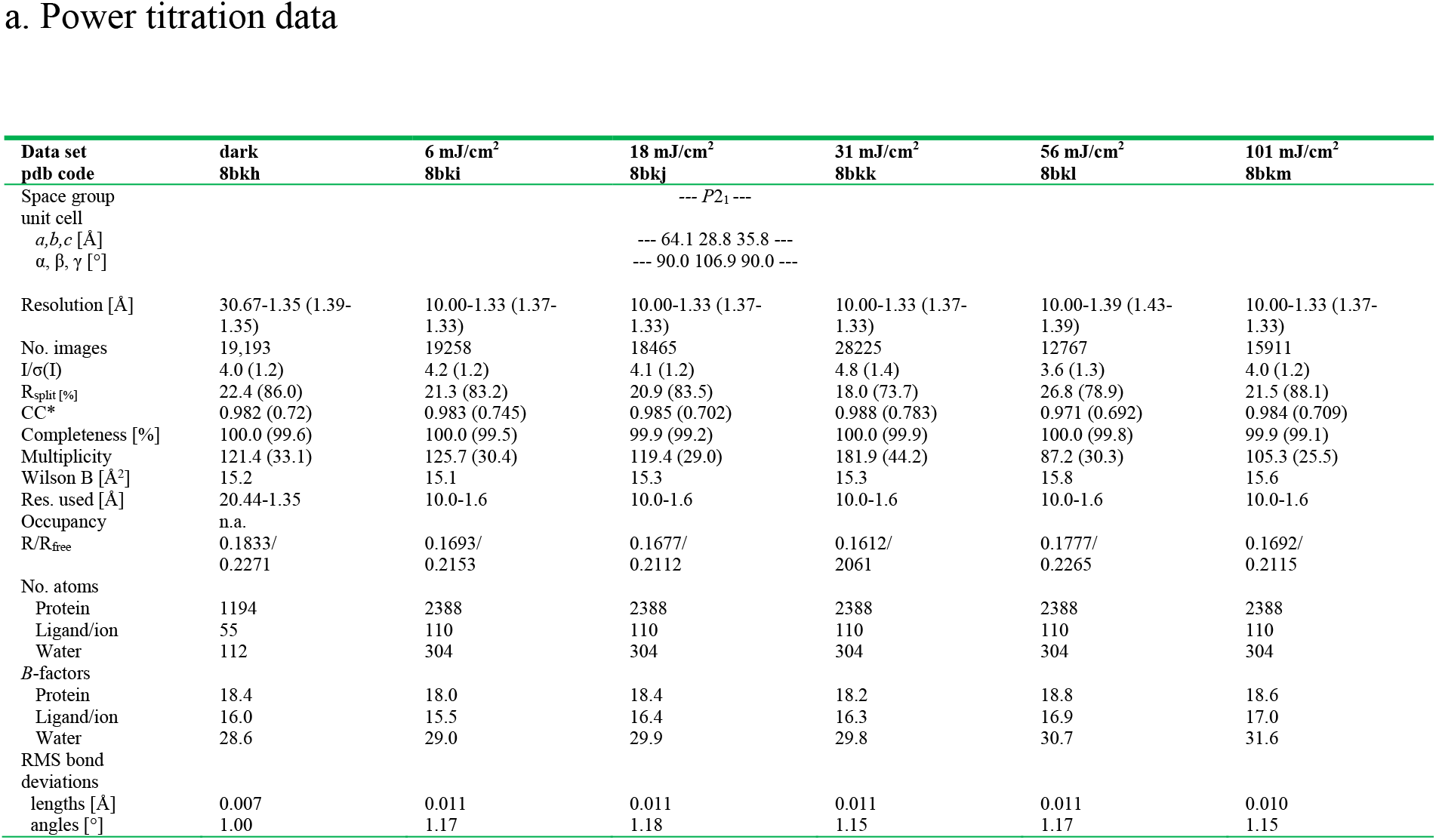
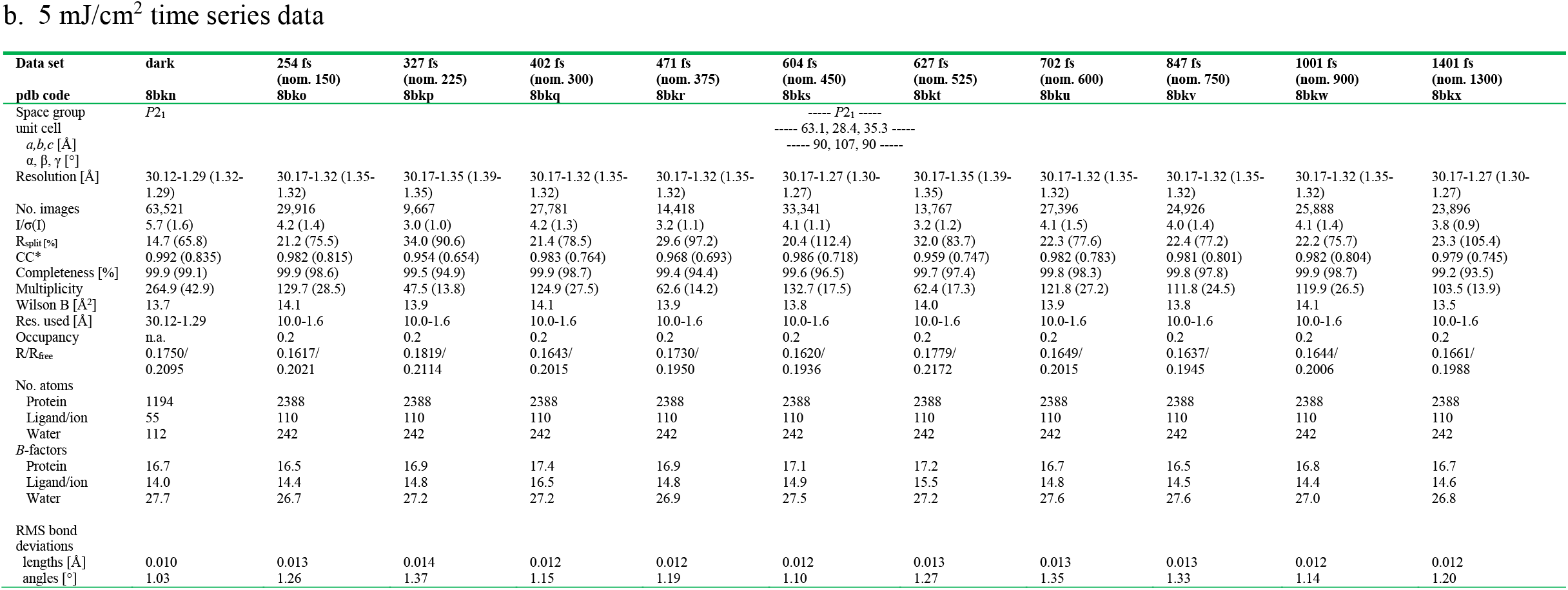
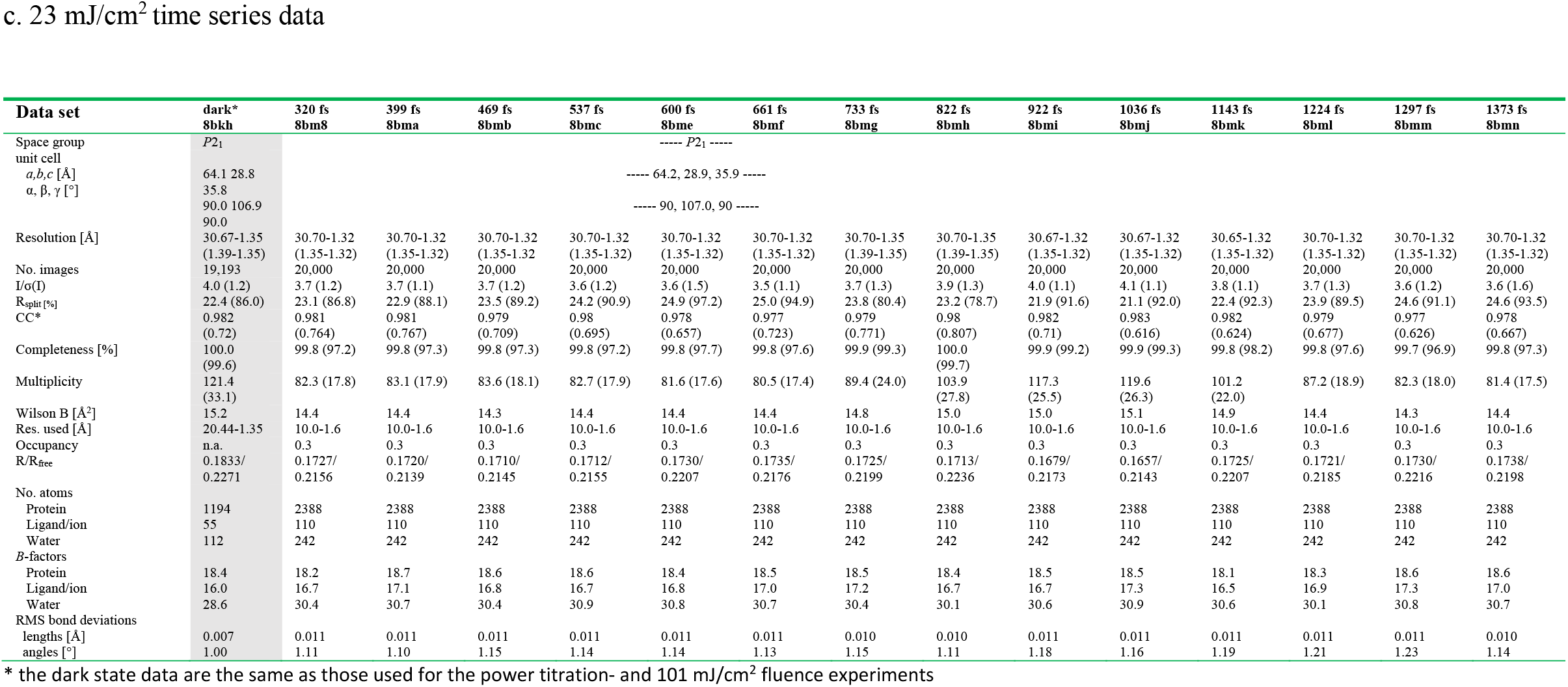
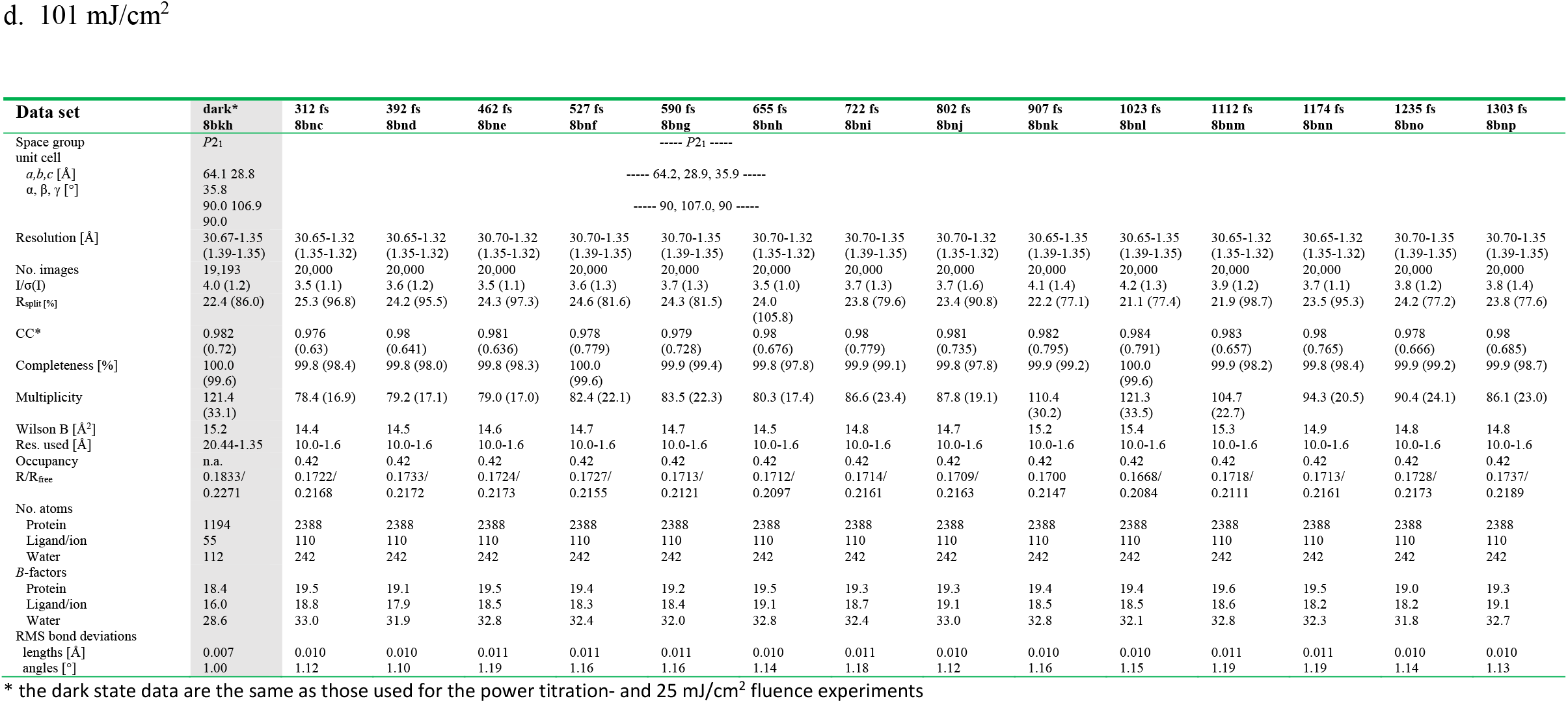
Crystallographic data and refinement statistics

**Extended Data Table2:**
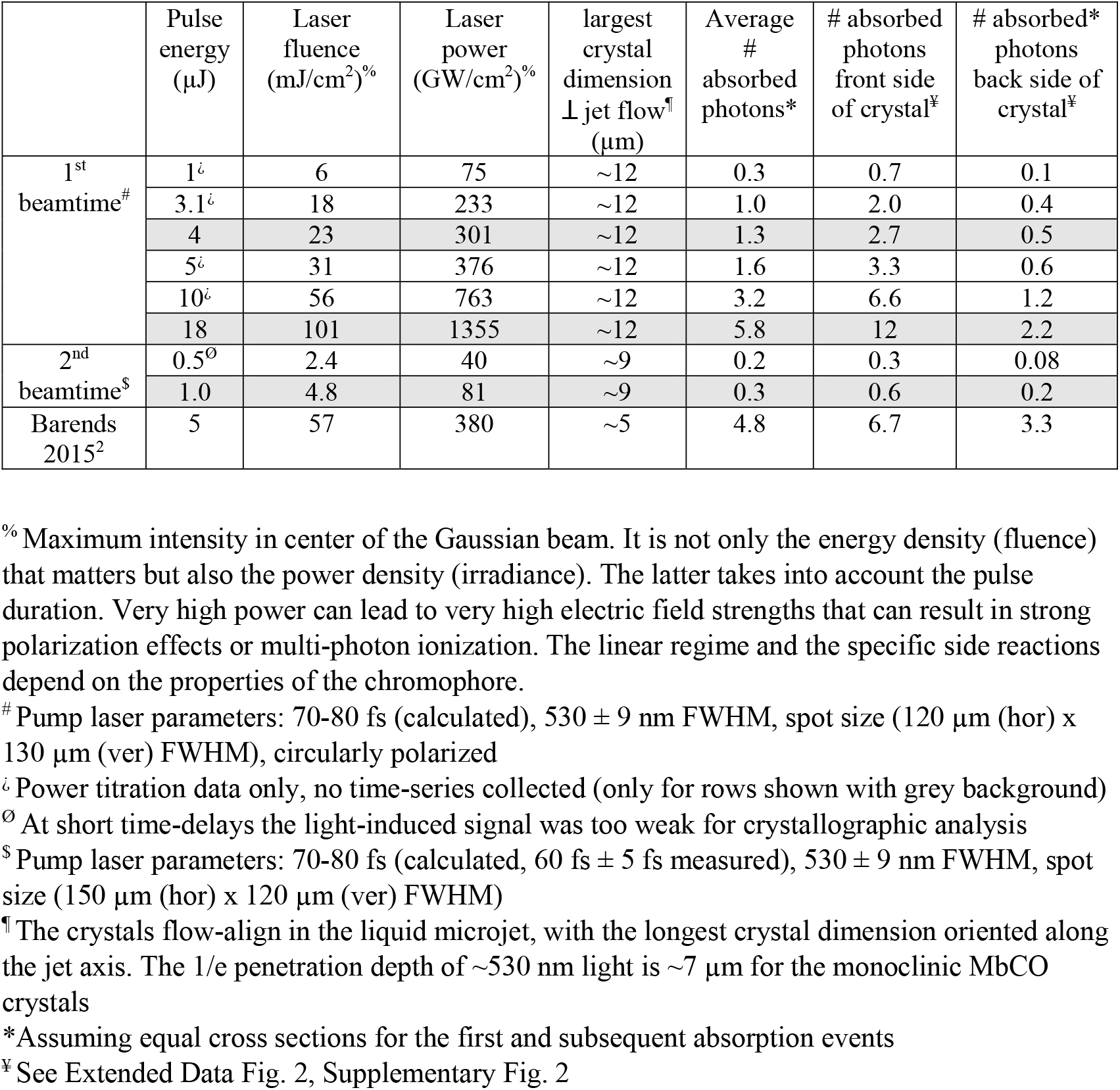
Laser and excitation parameters

**Extended data Figure 1.**
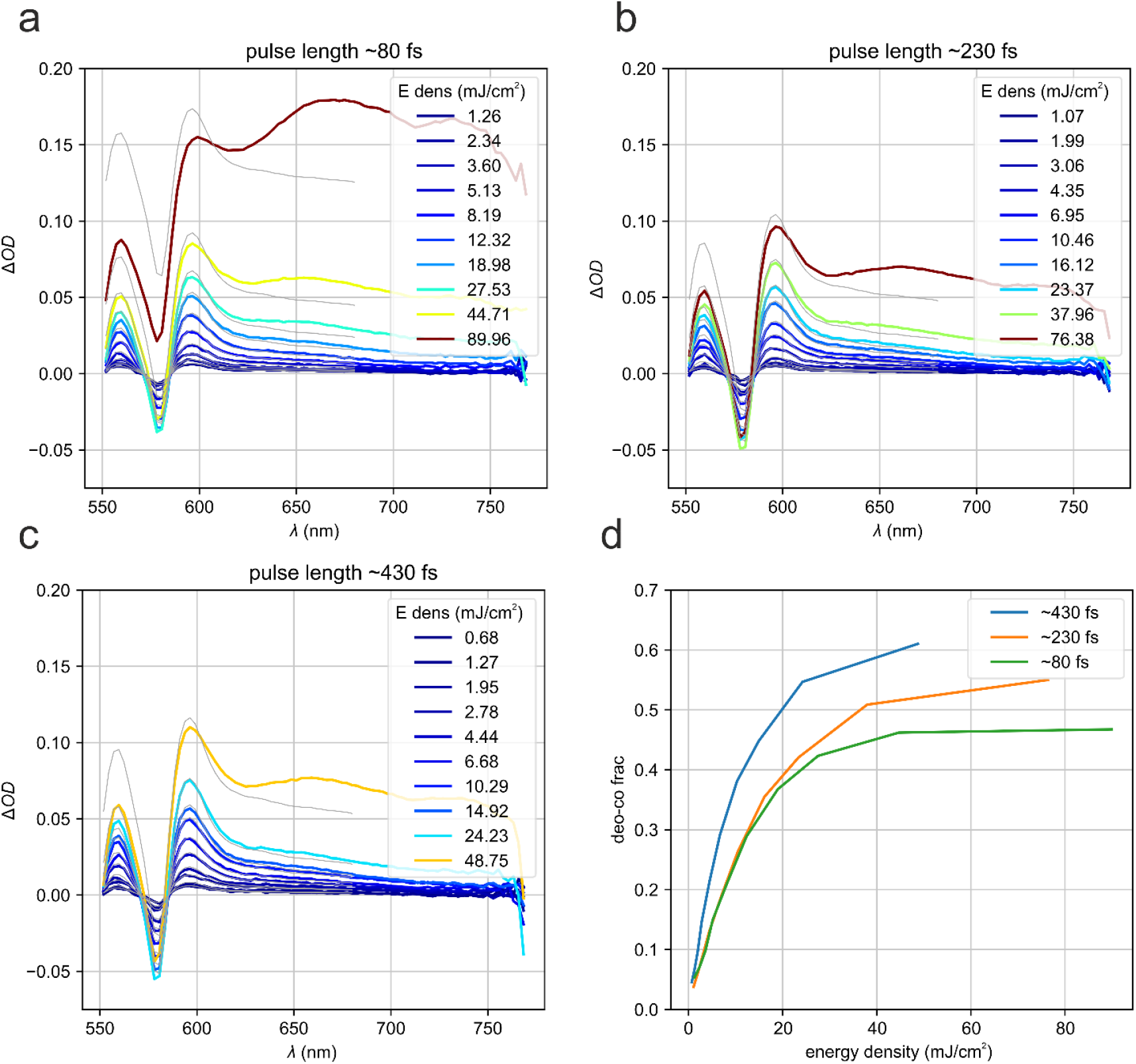
Optical power titration. Carboxymyoglobin solution (6 mM, 0.5 OD) was photoexcited using three different pump laser durations (∼80 fs (**a**), 230 fs (**b**), 430 fs, (**c**)) and different laser fluences, ranging from ∼0.7 mJ/cm^2^ to ∼90 mJ/cm^2^. Spectra were recorded after a 10 ps delay following a 532 nm laser pump pulse. The curves are color-coded with respect to the energy density (fluence, same color scale for a-b). The difference spectra, light-dark, were fit against difference spectra of deoxy myoglobin (the final state of the photodissociation reaction) and carboxymyoglobin (deoxyMb-MbCO + const. offset). The thin lines are the fits that were used to estimate the photolysis fraction shown in (d). At high laser intensity the spectra change shape and an additional peak appears around 650 nm with a lifetime of a few ps (data not shown). The longer pulses seem to yield a higher fraction of photoproduct. The plot shown in (d) allows identification of the linear photoexcitation regime; it is ≤10 mJ/cm^2^.

**Extended Data Fig. 2.**
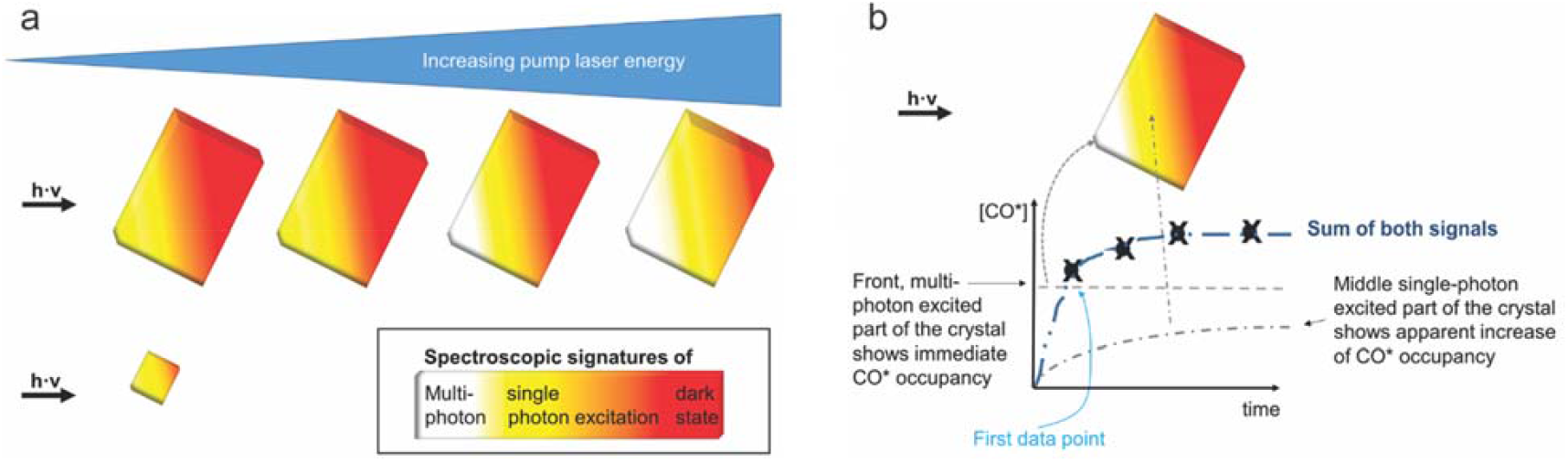
Crystal size, laser fluence and light-induced difference (light minus dark) signal are entangled quantities. a) Low intensity laser light, such as required for excitation in the linear excitation regime, cannot transverse crystals that have a dimension that exceeds the 1/e laser penetration depth. When this dimension is parallel to the laser beam, a significant fraction of the crystal volume cannot be photoexcited and a large pedestal of dark molecules remains (red), resulting in small light-dark differences. To increase the signal, the laser fluence is increased, which however results in multiphoton absorption at the front of the crystal. The issue is much reduced for crystals that have thickness d < 1/e of the pump laser penetration depth^19^. b) The different photoexcitation conditions in relatively thick crystals at high laser fluence can reflect onto the signal. For example, in the 101 mJ/cm^2^ fluence data the CO* signal increases strongly within the first time-delay, reflecting the very fast photodissociation upon multiphoton excitation at the “front” of the crystal. The increase of CO* with time is similar to the signal in the 5 mJ/cm^2^ and 23 mJ/cm^2^ data and originates from “deeper” regions in the crystal that were exposed to significantly lower fluence because of absorption by molecules in the “front”.

**Extended Data Fig 3.**
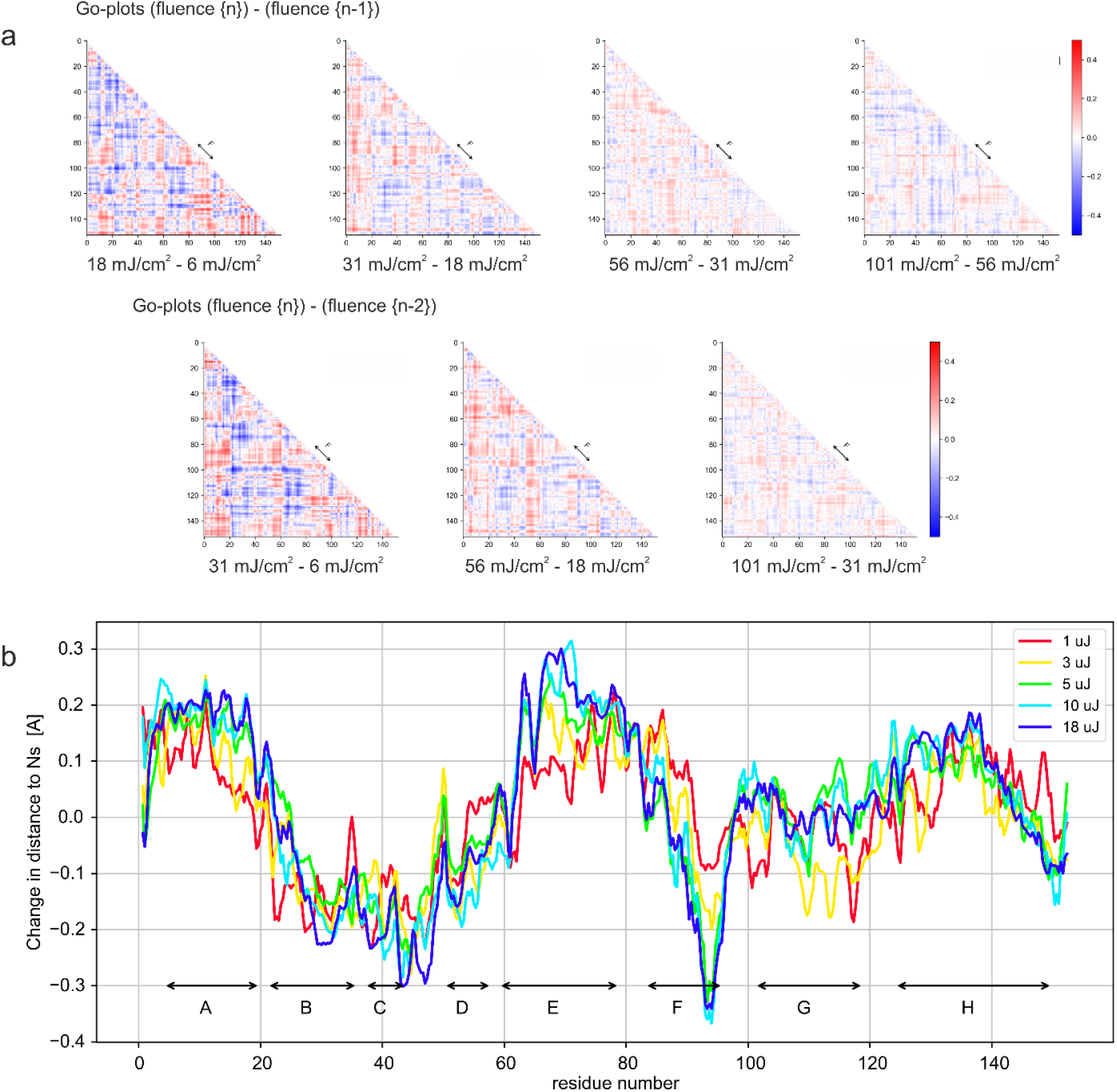
Structural changes as a function of pump laser fluence at a 10 ps time delay. To facilitate the identification of systematic differences between the structural changes observed upon photoexcitation at different laser fluences (see Fig. 1d), we calculated difference difference plots. The red and blue color-coding indicates that the atoms are further apart or closer together, respectively, than in the MbCO dark state structure. It appears that there are no systematic differences in correlated structural changes between pump laser energies. This differs from the findings described previously for a 3 ps time delay (see Fig. S13 in reference ^2^) showing a more pronounced displacement for example of the F-helix at a pump laser energy of 20 µJ (∼ 230 mJ/cm^2^; 1.5 TW/cm^2^) than at 6 µJ (∼70 mJ/cm^2^; 450 GW/cm^2^). In contrast, Guallar-type plots^29^, showing the change in distance of backbone N, Ca, and C atoms to the heme nitrogens for each time delay, show a clear difference for the different laser fluences.

**Extended Data Fig 4.**
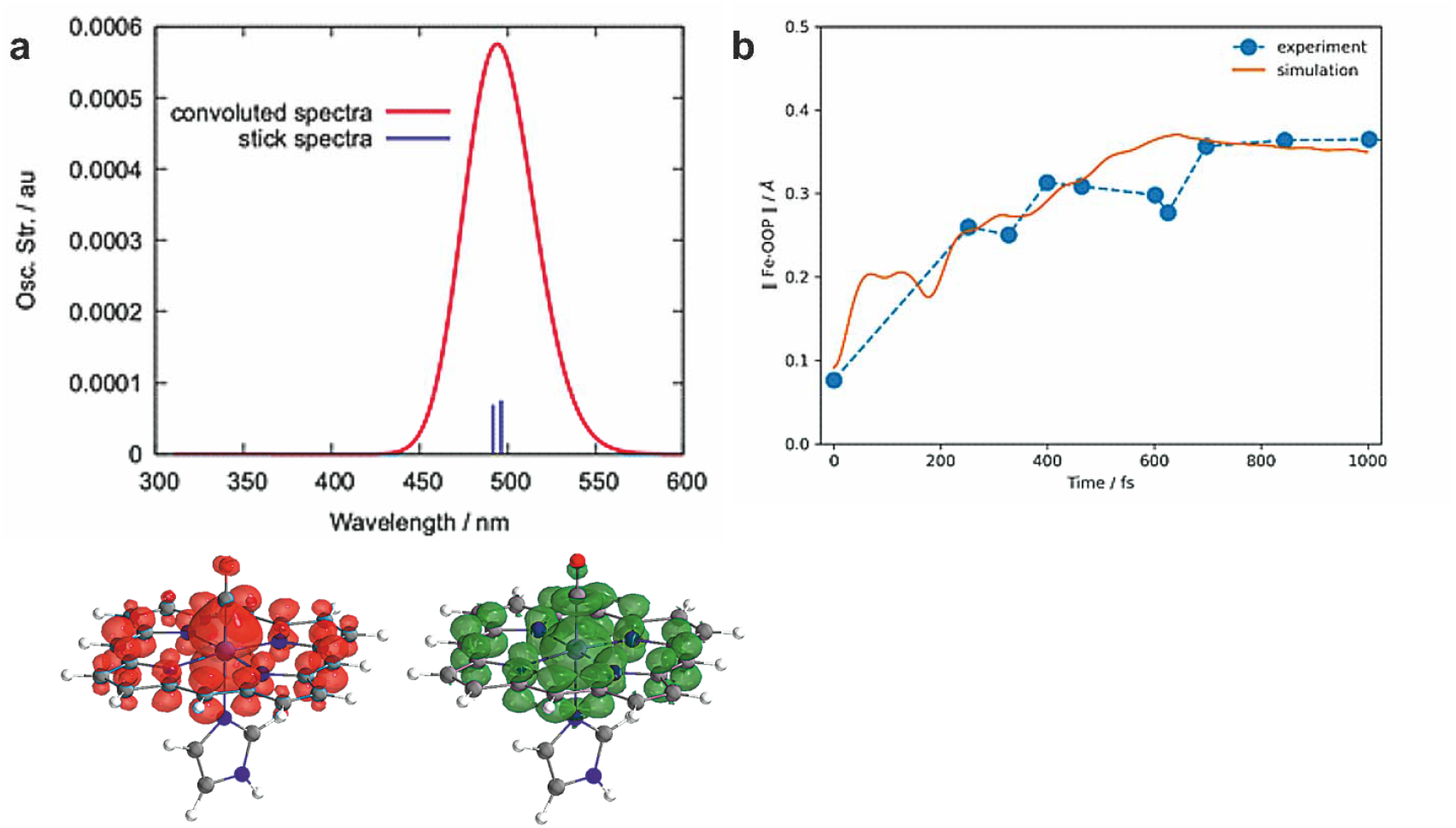
Quantum Chemistry simulations. a) The Q-band excited state absorption spectrum (top) can absorb to a high-energy singlet, in an excitation energy that is approximately two times that of the Q-band. This state corresponds to a mixed π→π* character of the heme and 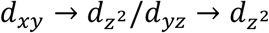 character with respect to the ground state, as analyzed from an attachment (green) / detachment (red) density analysis, showing that it is dissociative with respect to the Fe-CO bond. b) Comparison of the FeOOP motion derived by QM/MM dynamics (see Supplementary Note 2 for details) and TR-SFX (single photon excitation, 5 mJ/cm^2^ data).

**Extended data Fig 5.**
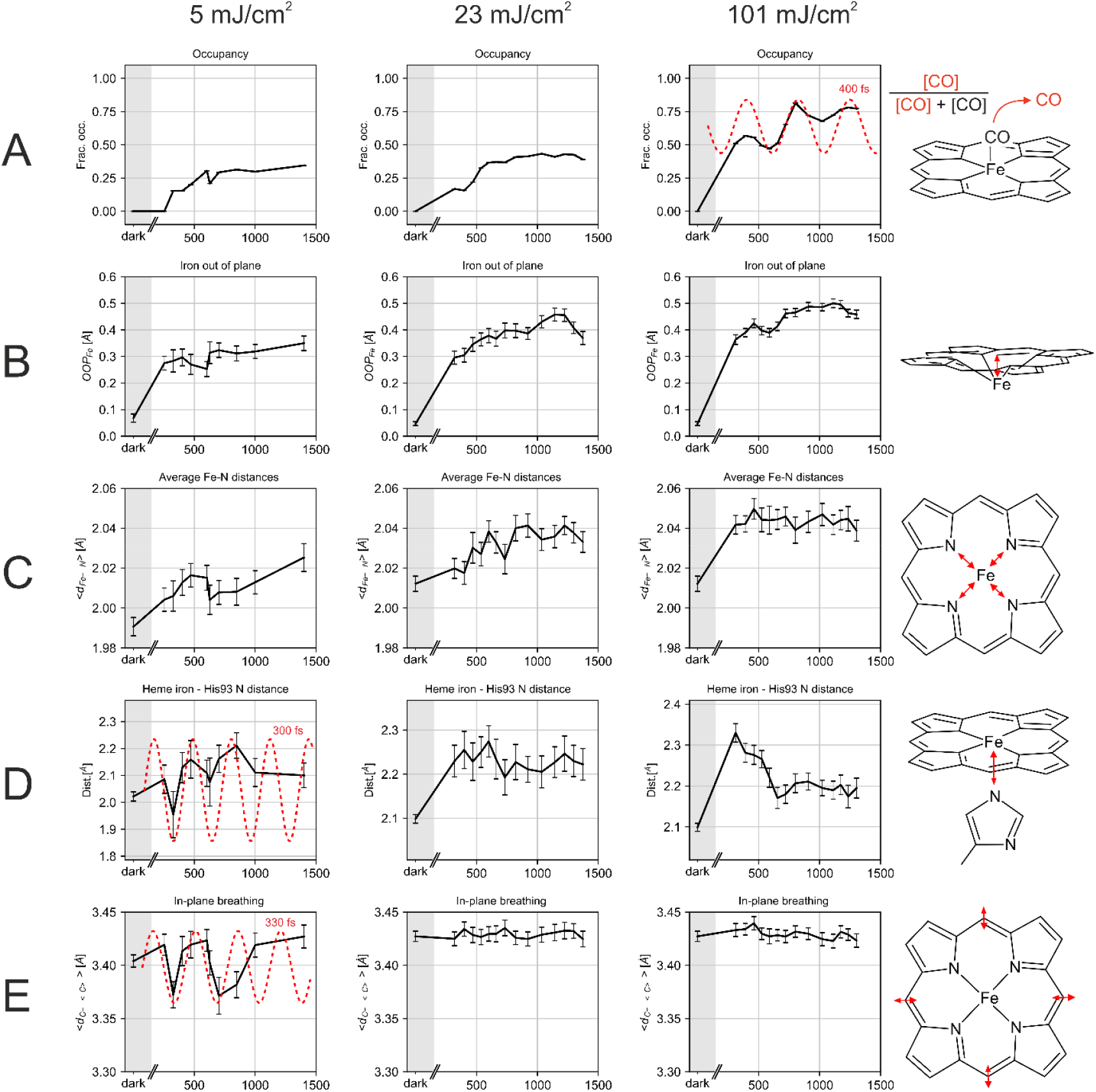
Heme structural dynamics. The figure corresponds to Fig. 2 in the main text but shows more details. a) Apparent CO* occupancy, check Fig. 2 legend for the temporal dependence of the 101 mJ/cm^2^ data. b) iron-out-of-plane distance, c) average distance between the iron atom and the porphyrin N atoms, d) distance between heme iron and proximal His93 NE2 atom, e) heme in-plane breathing (ν7 mode), determined as the average distance of the heme *meso* carbon atoms to the center of the heme. The oscillation periods are indicated by red dashed lines. The coordinate uncertainties are indicated; they were determined using bootstrapping resampling as described previously^48,60^, the error bars correspond to ±1 sigma.

**Extended Data Fig. 6.**
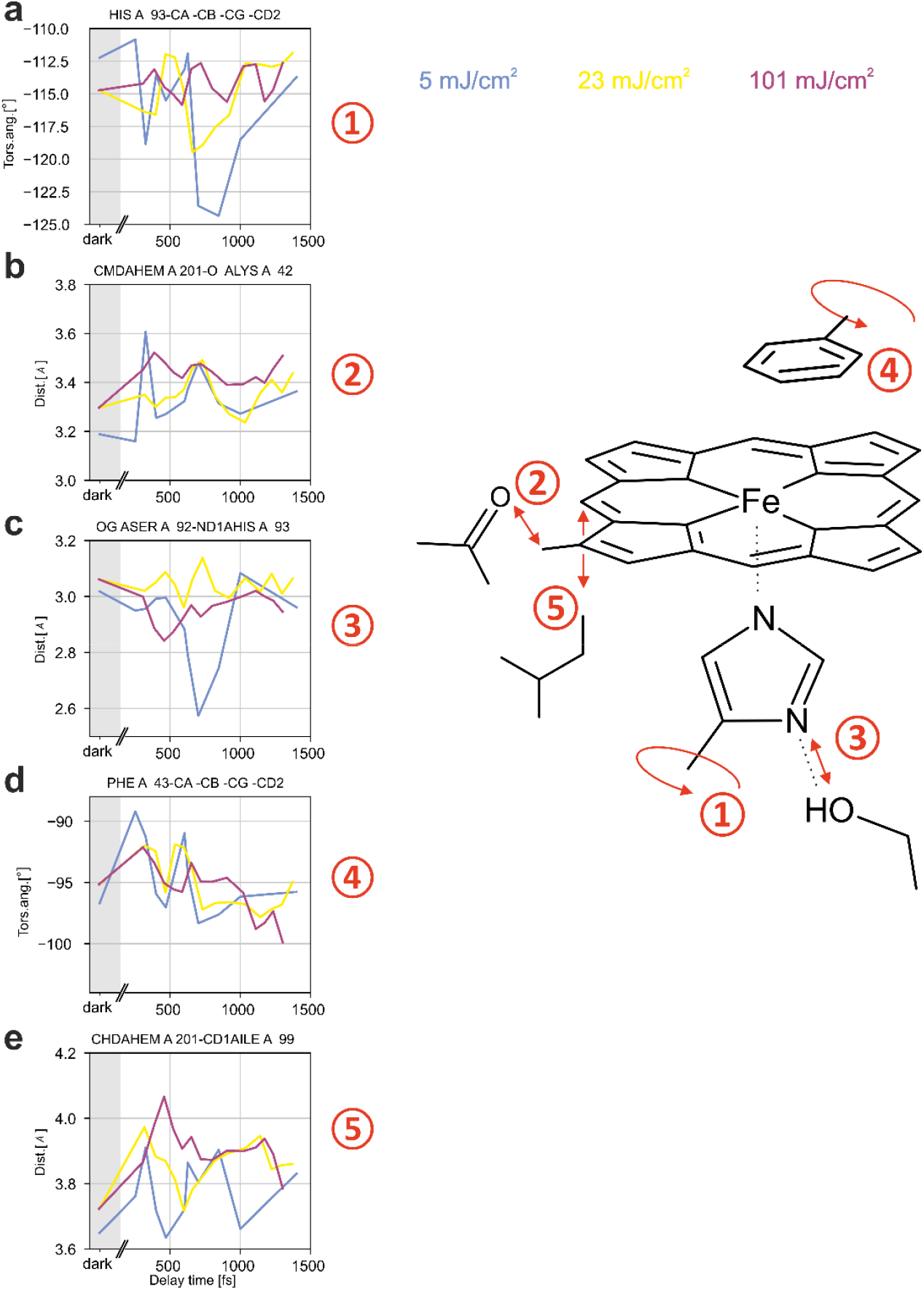
Dynamics of heme surroundings depend strongly on fluence. a) χ2 torsion angle of the heme-coordinating His93. At higher fluences, much smaller movements are observed than at 5 mJ/cm^2^. At 101 mJ/cm2 an oscillation is observed that is not apparent at lower fluences b) The distance between heme CMD atom and Lys42 backbone carbonyl O atom also shows different time evolutions with different fluences, with larger (and even oscillatory) motions at 5 mJ/cm^2^ but smaller motions at higher fluences Similar fluence-dependent effects are observed in c) the length of the His93 ND1…Ser92 OG hydrogen bond, d) the Phe43 χ2 torsion angle, and e) the heme CHD-Ile99 CD1 distance. Error bars corresponding to ±1 sigma and lines illustrating the oscillation periods are shown in Extended Data Fig. 7.

**Extended Data Fig. 7.**
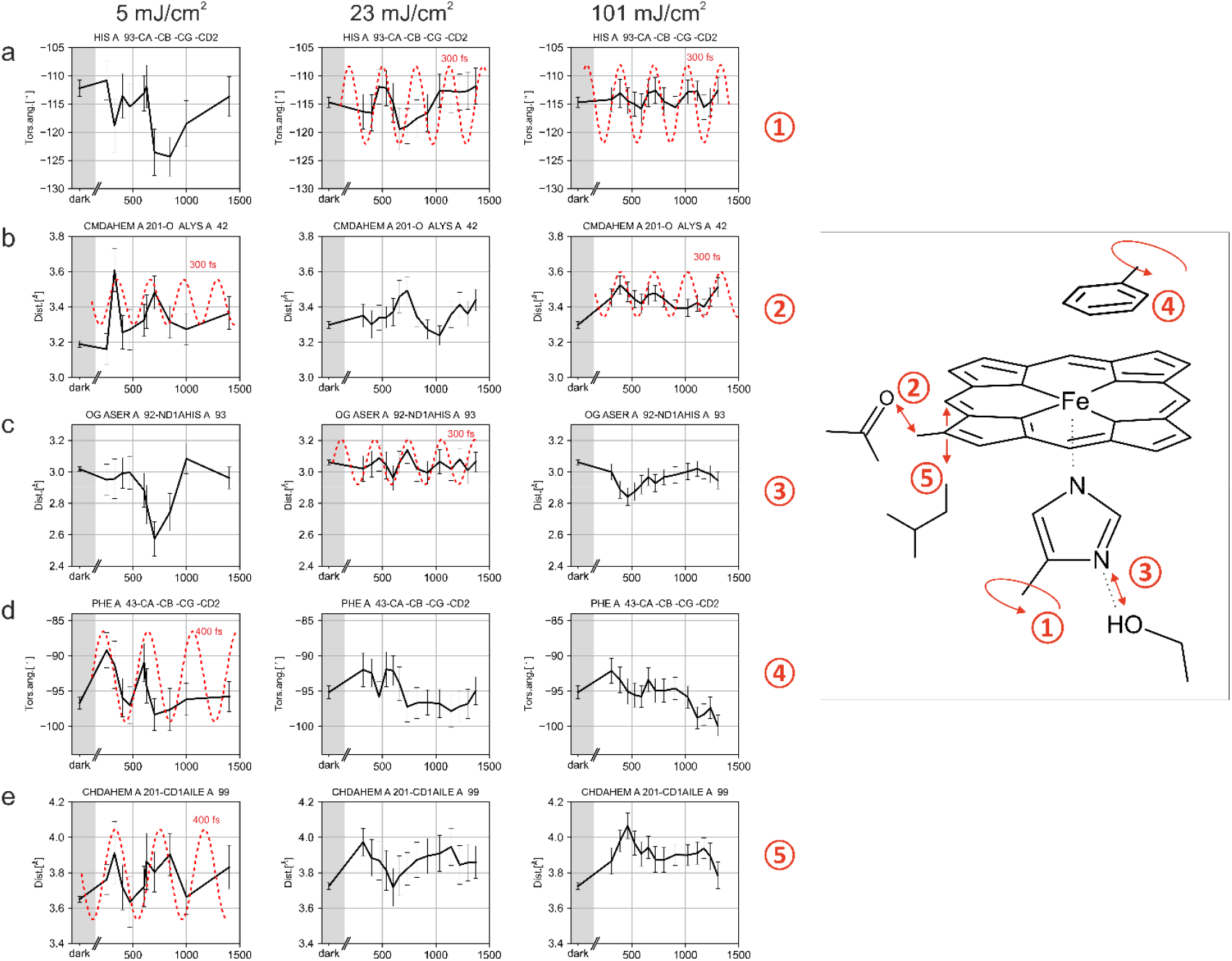
Dynamics of heme surroundings. a) His93 χ2 torsion angle, b) Distance between heme CMD atom and Lys42 backbone carbonyl O atom, c) Length of the His93 ND1…Ser92 OG hydrogen bond. d) Phe43 χ2 torsion angle, e) heme CHD-Ile99 CD1 distance. Red dashed lines illustrate oscillation periods. The coordinate uncertainties are indicated; they were determined using bootstrapping resampling as described previously^48,60^, the error bars correspond to ±1 sigma.

**Extended Data Fig. 8.**
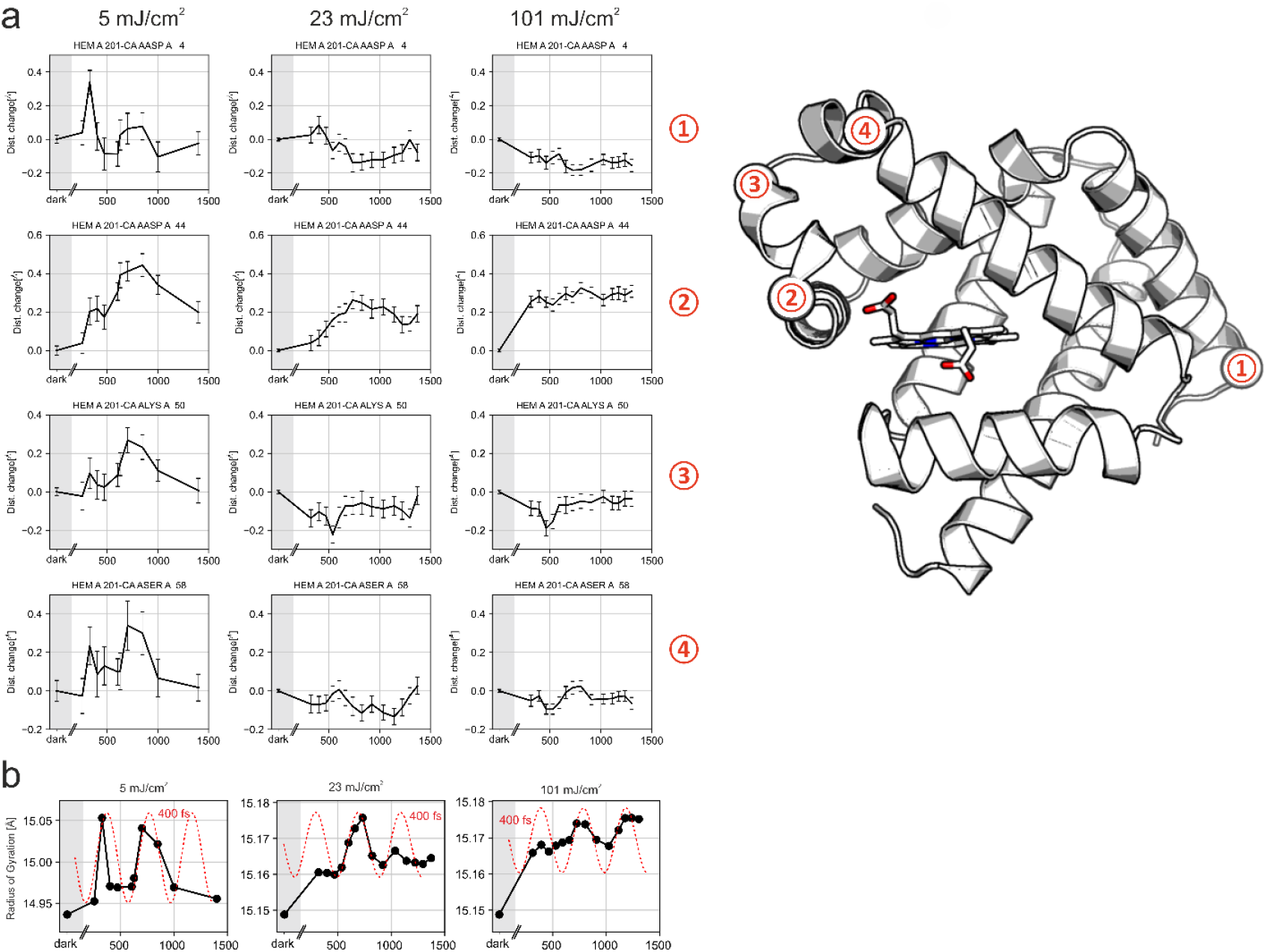
C*α* atoms at the end of helices show an oscillatory modulation with time at low photoexcitation fluence. Left panel: 5 mJ/cm^2^, middle panel 23 mJ/cm^2^, right panel 101 mJ/cm^2^. The location of the residues, chosen to be at the beginning or end of helices, is indicated. The F-helix (located below the heme, parallel to its plane) is shown in Fig. 3. The oscillatory modulation of the structural changes is also apparent in the temporal evolution of the radius of gyration Rg. Also in this case, the oscillation is strongest and most pronounced in the 5 mJ/cm^2^ data, corresponding to photoexcitation in the linear regime. Red dashed lines illustrate oscillatory periodicities. The coordinate uncertainties are indicated; they were determined using bootstrapping resampling as described previously^48,60^, the error bars correspond to ±1 sigma.

