## Supplemental information for "Influence of pump laser fluence on ultrafast structural changes in myoglobin"

**Supplementary Information, Barends et al.**

Supplementary Notes 1,2

Supplementary Figures 1-4

**Supplementary Note 1:** TR-SFX pump probe experiments face conflicting demands with respect to crystal dimensions: high-resolution diffraction typically benefits from thick crystals whereas efficient photoexcitation requires optically thin samples. Specifically, absorption as described by Lambert-Beer's law sets a strict upper limit for light penetration of a given wavelength, absorption coefficient and chromophore concentration. Due to the high chromophore concentration in crystalline samples (typically 10-30 mM), this in turn translates into a need for very small crystal dimensions. One can change the pump wavelength to reduce absorption and thereby allow use of larger crystals, but only while bearing in mind any possible concomitant changes in cross sections of the various pathways. These often reduce the yield of the desired photoproduct[1,2] which, after all, is ultimately the quantity to be optimized. Thus, experimental conditions need to be identified such that maximal population inversion is obtained in the probed crystal volume upon excitation with a femtosecond laser pulse, while yet ensuring that the photoproduct yield is intrinsically limited only by internal conversion reactions[1].

In pump probe TR-SFX experiments a quick estimate of the photoproduct yield – or that of an ensuing reaction intermediate – can be obtained by calculating $F_{obs}^{light}$- $F_{obs}^{dark}$ difference electron density maps and plotting the height of the positive and negative peaks (corresponding to appearance and disappearance of light and dark state, respectively) as a function of laser energy (see Fig. 1a,b). If the signal is inadequate, the laser energy is then increased. Another pump probe SFX dataset is collected and new difference electron density maps calculated. The objective, however, must not be to simply *increase* peak heights until usable signal results. Valid characterization of biologically relevant single photon-induced reactions can only be made in the regime of *linearly increasing* signal with pump laser energy. Establishing the linear regime requires experimental power titrations. Nonlinearities observed at high pump fluences can arise due to various factors (absorption saturation leading to side reactions triggered upon multiphoton absorption (stimulated emission, photoionization, …). Consequently, power density is critical, meaning that both energy density and pulse length of the pump laser play a role.

If the crystal size exceeds the 1/e penetration depth of the pump laser wavelength, high laser fluence often seems unavoidable in obtaining adequate photoexcitation. This is because very few photons reach the “rear” regions of the crystals, forming a “pedestal” of dark molecules within each crystal (Extended Data Fig. 2). The resulting small apparent $F_{obs}^{light}$- $F_{obs}^{dark}$ difference is simply due to inadequate excitation of chromophores in this pedestal at low laser fluence. Note also that it is impossible to adequately excite these “rear” molecules without also overexposing the “front” molecules, resulting in their multiphoton excitation. Often it is assumed that this is not an issue for thin plate-shaped crystals. For randomly oriented crystals (as in SFX) this is incorrect.
A frequent justification for using relatively large crystals for time-resolved experiments is that smaller crystals often do not diffract to high resolution, in particular crystals of very anisotropic dimensions (needles, plates). However this issue can be addressed by fracturing larger crystals into microcrystalline parts[3].

There are several reasons that the light-induced signal can be much lower than expected: (i) losses due to scattering, for example at air-liquid interfaces, resulting in lower than expected laser fluence at the crystal, (ii) unfavorable orientation of the absorption dipole moment of the chromophore **μ** and the electric field vector **E**. The probability of excitation is proportional to $(\boldsymbol{\mu}\cdot\mathbf{E})^2$ and thus ~$\cos^2 \theta$, with θ being the angle of the laser polarization vector and the chromophore transition dipole. This effect is particularly noticeable at low laser fluence, and in crystals significantly more so than in solution. Since the orientations of chromophores within a crystal are anything but stochastic, it can even be that for certain crystal orientations no photons at all are absorbed. (iii) stimulated emission (SE). Whether a photon is absorbed or whether it stimulates emission depends on the ground and excited state populations (the Einstein coefficients for absorption and SE are the same). Loss through SE dominates at high laser fluence, see for example reference [4].

For jet-based measurements, scattering losses through (i) are typically ≤20 % for non-scattering carrier media such as LCP and other transparent media. However any optically small scattering centers (small water droplets in grease[5] or other tiny particles in the crystal-surrounding medium, notably the particles intrinsic to Super Lube[6]) result in strong isotropic scattering of the pump light within the jet. This makes quantification of the effective laser irradiance at the crystal extremely difficult. Crystals outside of the direct light path can be illuminated, as can crystals upstream of the illumination zone[7]. Simple transmission measurements then yield misleading effective fluences[5]. A general problem when reporting scattering losses at jet interfaces is the neglect of the strong lens effect of the cylindrical surface of the jet[8], resulting in erroneously high reported losses of ~ 80 %[9].

**Supplementary Note 2. Quantum chemistry.**

The initial steps of the MbCO photolysis are now well established[10]. Absorption of a photon to the singlet Q state (Supplementary Fig. 3a) is followed by transfer to the singlet metal-to-ligand charge-transfer (MLCT) band due to a strong Jahn-Teller distortion in the excited state. This affords an energy transfer from the porphyrin plane to the Fe-center, activating dissociative Fe-CO stretching vibrations and thus CO dissociation (see Supplementary Fig. 3b). Sequential Fe spin transitions ensue, ultimately resulting in the formation of a high-spin FeII center.

We showed recently that the Jahn-Teller distortion ultimately induces coherent oscillations of the Fe-CO bond distance[10]. The predicted damping time is in line with the apparent increase of the occupancy of photolyzed CO with time (Fig. 2a)), interpreted as decreasing “disorder” of the position of the dissociated CO with time. However, with increasing photon flux, the apparent heme-CO photolysis appears “accelerated” and, eventually, the apparent occupancy of CO hardly changes with time (Fig. 2a, ref. [11]). We explain this observation by sequential two-photon absorption. The first photon leads to the Q-band excitation of heme-CO (Supplementary Fig. 3a) corresponding to a $\pi \rightarrow \pi^*$ transition in porphyrin. This state can absorb a second photon. The absorption spectrum of the Q-excited heme-CO system is shown in Supplementary Fig. 3c, in which a high energy single state is populated, corresponding to roughly two times the energy of the Q state. Analysis of its excitation character shows that this state is mixed $\pi \rightarrow \pi^*$ of the heme and $d_{xy} \rightarrow d_{z2}$/ $d_{yz} \rightarrow d_{z2}$ character with respect to the ground state, and therefore is dissociative for the Fe-CO bond (see Supplementary Fig. 3d).

We performed a relaxed scan of the potential-energy surface (PES) cut of the singlet manifold along the Fe-C(O) dissociation coordinates (see Methods, Computational Details section). Supplementary Fig. 3e depicts the lowest 60 electronic states in grey. The black lines represent the

Q states, the red line represents the higher-energy dissociative state (Supplementary Fig. 3e). It is clear that upon excitation to the dissociative singlet, the excited wave packet experiences a large gradient towards Fe-C(O) dissociation and due to the (barrierless) repulsive nature of the potential, no coherent oscillation of the wave packet is expected on this state.

The Fe out-of-plane (OOP) motion is one of the quantities that is measurable from the experimental data during the MbCO photolysis. The dynamics of myoglobin following photodissociation of CO has been simulated at the QM/MM level. It can be assumed that the difference in the energies of the bright states and the lowest $^5$MLCT state is deposited as kinetic energy to the atoms closest to the chromophore. The amount of excess energy (0.1 eV) is redistributed randomly to the kinetic energy of the QM atoms of the simulation box. From this, 40 initial conditions were generated and propagated up to 1 ps at the DFT/BHHLYP/6-31G* level. The ensemble average of the Fe-OOP coordinate was computed as a function of time. The results of the simulation agree well with the experimentally derived ones (see Supplementary Fig. 3f). The Fe-OOP coordinate starts from its equilibrium value (approximately 0.1 Ang.) in MbCO and increases with time up to shortly after 600 fs. The Fe-OOP coordinate does not change significantly after 600 fs both in experimental and simulation results.

In order to interpret the dynamics at different time delays, we analyzed the protein cavity changes during a relaxed scan coordinate from a sudden dissociation of CO (Supplementary Fig. 4). The reaction coordinate (Supplementary Fig. 4a) corresponds to the different Fe-CO distances.

The scans show conformational changes in both the heme (Supplementary Fig. 4b-e) and the surrounding protein (Supplementary Fig. 4f-j) along this reaction coordinate, which in several cases show the same trends as observed by time-resolved SFX (Extended Data Fig. 5 and 7), such as for the heme CHD-Ile99 CD1 distance, the Phe43 $\chi 2$ torsion angle and the iron out-of-plane motion.

**Supplementary Figures**

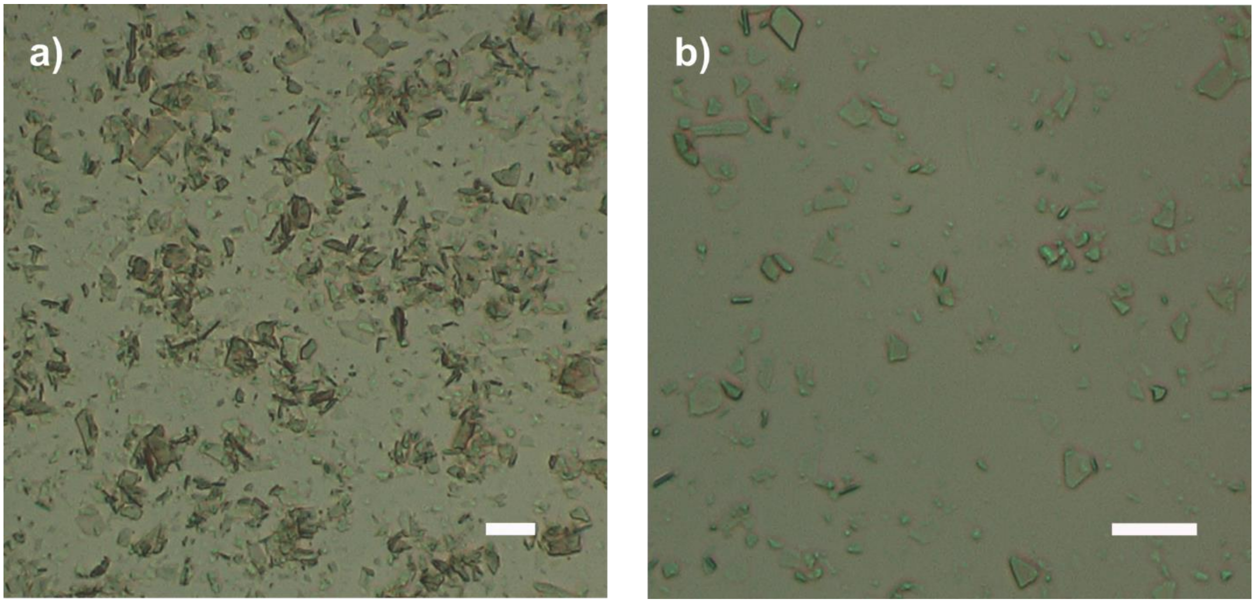


**Supplementary Figure 1**

Macroscopic myoglobin crystals were fractured by filtration using tandem array stainless steel ¼ inch diameter filters[3]. For the first beamtime two filter stacks were used consisting of 100, 40 µm filters followed by a second tandem array of 40, 20, 10, 10, 5 µm. This resulted in the crystallites shown in (a), with many of the larger crystals showing lengths of ~ 15-20 µm. For the second beamtime the crystalline slurry was further fractionated using a tandem array of 10, 5, 2, 2 µm filters. The larger crystals are less than 10 µm long. The white scale bar corresponds to 20 µm. Images were taken with a Hirox microscope.

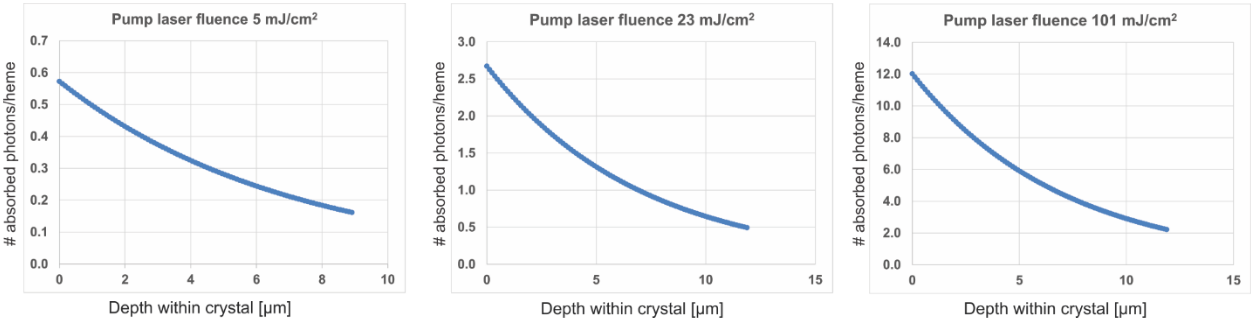


**Supplementary Figure 2. Average number of nominally absorbed photons.** The number of nominally absorbed photons/heme in photoexcited monoclinic myoglobin crystals was calculated for different pump laser energies as described recently[8]. It was assumed that the first and subsequently absorbed photons have the same absorption cross sections. As indicated (see Extended Data Table 2), the crystals used for SFX data collection of the 5 mJ/cm$^2$ laser fluence data were smaller than those used for the higher fluence photoexcitation data.

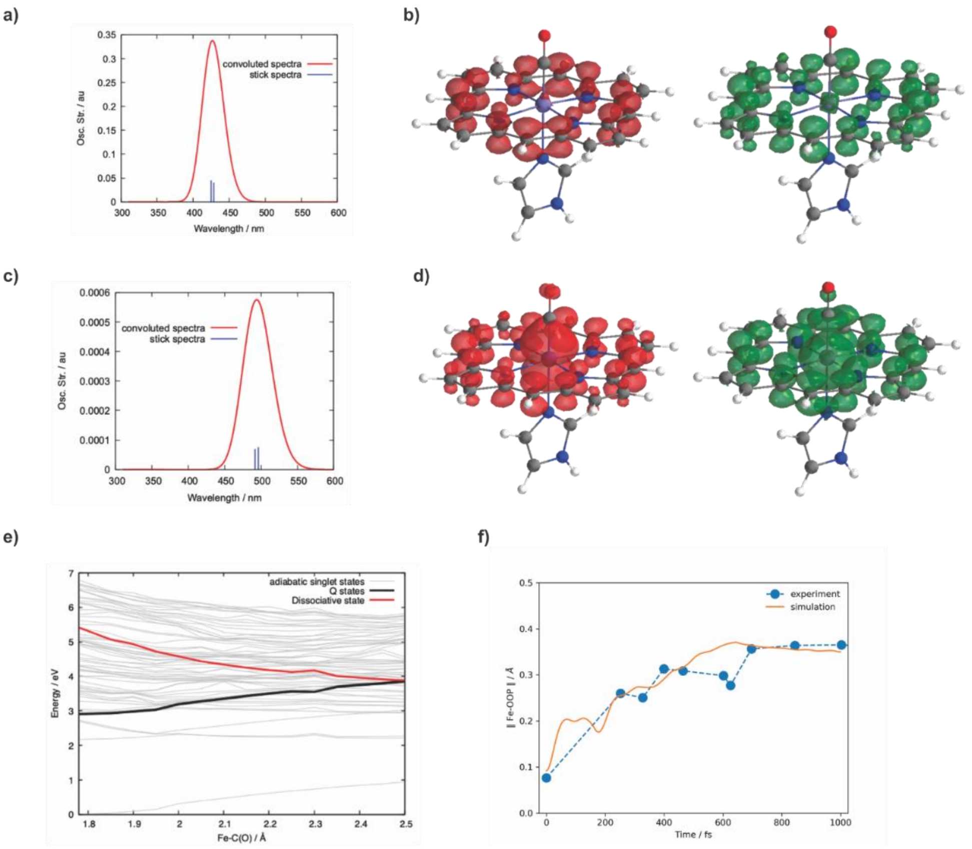


**Supplementary Figure 3**: a) Single photon absorption. Absorption spectrum of the heme-CO model ground state; b) Attachment-detachment density analysis of the most intense transition in this spectral region, corresponding to the Q states of porphyrin. In red, the electron density is depleted and in green the electron density is increased. c) Sequential two-photon absorption. Absorption spectrum of the Q-band excited heme-CO model. d) Attachment-detachment density analysis of the most intense transition in this spectral region, corresponding to a dissociative Fe-CO state. In red, the electron density is depleted and in green the electron density is increased. e) Potential energy curve of 60 electronic singlet states along a relaxed scan on the Fe-C(O) dissociation. The grey thin lines represent the adiabatic states. A diabatic character is represented in thicker colored lines; the black thick lines represent the Q state and the red thick line depicts the dissociative state. f) cumulative running average (orange line) of the Fe-OOP coordinate as a function of time for the 40 QM/MM trajectories overlaid with the single photon excitation experimental data (5 mJ/cm$^2$) (blue points).

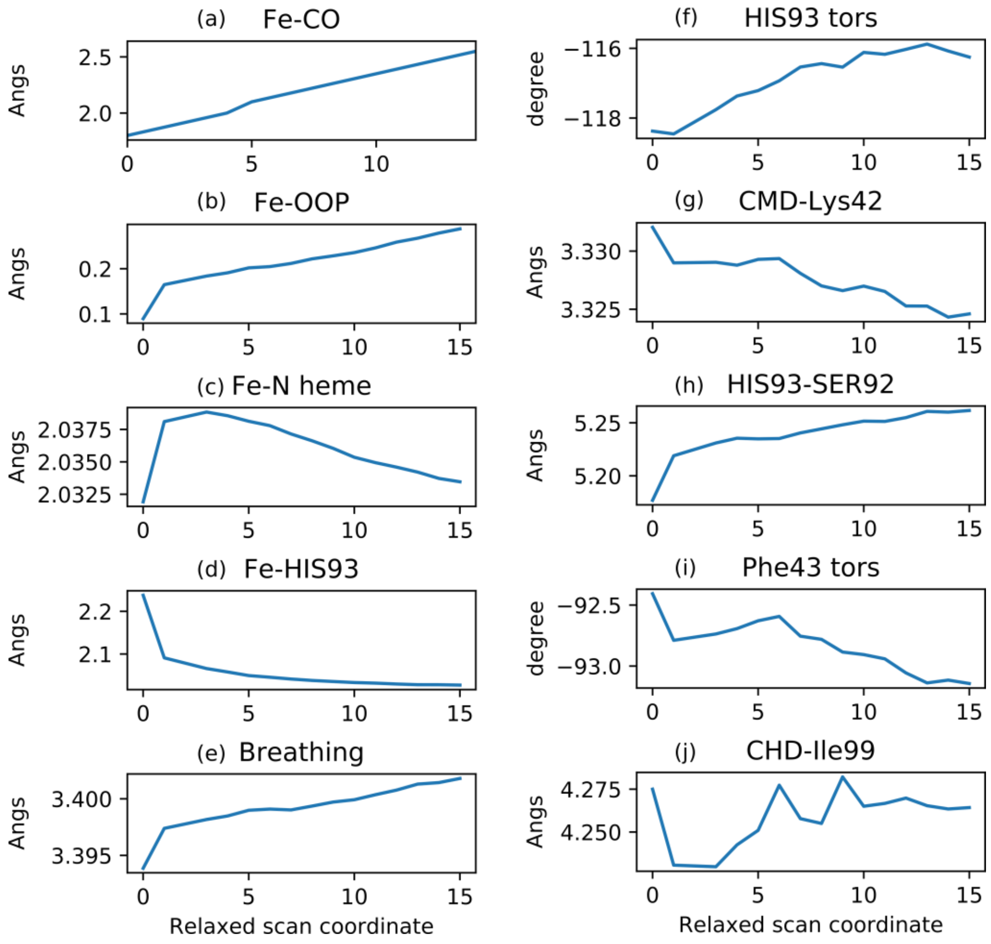


**Supplementary Fig. 4**. Changes in the heme and protein cavity motions along a relaxed scan coordinate in the protein model. a) definition of the relaxed scan coordinate at fixed Fe-CO distances. b) Iron out-of-plane (FeOOP) motion. c) Average iron-nitrogen (Fe-Np) distance. d) Iron-nitrogen distance from the proximal histidine His93 (Fe-$N_{His93}$). e) Heme in-plane breathing mode. f) His93 $\chi$2 torsion angle, g) distance between the heme CMD atom and the Lys32 backbone carbonyl atom, h) length of the His93 ND1…Ser92OG hydrogen bond, i) Phe43 $\chi$2 torsion angle, j) heme CHD-Ile99-CD1 distance.
